# Spatiotemporal Control of Genomics and Epigenomics by Ultrasound

**DOI:** 10.1101/2023.06.21.544125

**Authors:** Yiqian Wu, Ziliang Huang, Yahan Liu, Chi Woo Yoon, Kun Sun, Yinglin Situ, Phuong Ho, Zhou Yuan, Linshan Zhu, Justin Eyquem, Yunde Zhao, Thomas Liu, Gabriel A Kwong, Shu Chien, Yingxiao Wang

**Author notes:** These authors contributed equally (Yiqian Wu, Ziliang Huang, Yahan Liu). Corresponding authors, (Y. Wang); (Y. Wu).

## Abstract

CRISPR (clustered regularly interspaced short palindromic repeats) is a revolutionary technology for genome editing. Its derived technologies such as CRISPR activation (CRISPRa) and CRISPR interference (CRISPRi) further allow transcriptional and epigenetic modulations. Focused ultrasound (FUS) can penetrate deep in biological tissues and induce mild hyperthermia in a confined region to activate heat-sensitive genes. Here we engineer a set of CRISPR(a/i) tools containing heat-sensitive genetic modules controllable by FUS for the regulation of genome and epigenome in live cells and animals. We demonstrated the capabilities of FUS-inducible CRISPRa, CRISPRi, and CRISPR (FUS-CRISPR(a/i)) to upregulate, repress, and knockout exogenous and/or endogenous genes, respectively, in different cell types. We further targeted FUS-CRISPR to telomeres in tumor cells to induce telomere disruption, inhibiting tumor growth and enhancing tumor susceptibility to killing by chimeric antigen receptor (CAR)-T cells. FUS-CRISPR-mediated telomere disruption for tumor priming combined with CAR-T therapy demonstrated synergistic therapeutic effects in xenograft mouse models. The FUS-CRISPR(a/i) toolbox allows the remote, noninvasive, and spatiotemporal control of genomic and epigenomic reprogramming in vivo, with extended applications in cancer treatment.

---

The emergence of CRISPR technology has revolutionized numerous aspects of life science and medicine^1–5^. With a single guide RNA (sgRNA), the Cas9 nuclease can be targeted to, in principle, any accessible genomic locus next to a protospacer adjacent motif (PAM) to cause site-specific double-strand break (DSB), providing a powerful way of editing endogenous genome and ultimately the phenotypes of organisms^6, 7^. The subsequent development of CRISPRa and CRISPRi with nuclease-dead Cas9 (dCas9) further enabled transcriptional and epigenetic modifications of endogenous loci, demonstrating the power of CRISPR in regulating the genome at different levels^8, 9^. As the CRISPR-based technologies advanced to translational applications and clinical trials, safety/controllability has become one of the major concerns, mainly due to the immunogenicity of Cas9-related proteins and their off-target effects accumulated during long-time expression in the cells^10–12^.

To address this, controllable CRISPR systems utilizing small molecules^13–15^, light^16–19^, or heat^20, 21^ as external cues for induction have been developed. Small-molecule-based systems can tightly control the time of action for CRISPR, but the diffusive characteristic of small molecules compromises the spatial precision. Light-based systems provide an elegant solution to control both the timing and location of CRISPR; however, it requires light-sensitive proteins which can be bulky and difficult to deliver, or possibly immunogenic due to their non-human origins^22, 23^. Also, the penetration depth of light with a maximum of millimeters limits its therapeutic applications particularly in tissues tens of centimeters deep^24^. The previously reported heat-inducible CRISPR-dCas9 systems rely on near infrared light stimulation aided by intermediate nanorods, which once again suffers from limited controlling depth in vivo^20, 21^.

Focused ultrasound (FUS) can penetrate deep and directly induce localized hyperthermia without intermediate co-factors in biological tissues^25, 26^. In fact, it has been used for tissue ablation in patients at relatively high temperatures (>60 °C)^27–30^, and for controlling heat-sensitive transgene expression in vivo at mildly elevated temperatures (42 - 43 °C)^31–36^. We have previously developed FUS-inducible CAR (FUS-CAR)-T cells that can be acoustogenetically activated by FUS for cancer therapy with reduced off-tumour toxicities^37^. The penetration power of FUS and its spatiotemporal precision allow the direct control of CRISPR without co-factors for genome editing and regulations at specific tissues and organs.

Here, we have developed a set of acoustogenetics- and CRISPR-based tools that include FUS-inducible CRISPRa (FUS-CRISPRa), FUS-inducible CRISPRi (FUS-CRISPRi), and FUS-inducible CRISPR (FUS-CRISPR). We have shown that this FUS-CRISPR(a/i) toolbox can allow FUS-inducible genomic and epigenomic reprogramming in multiple cell types and in vivo for synergistic therapeutics.

## Results

FUS can generate localized and mild hyperthermia in biological tissues. The heat stress can be sensed by cells through the endogenous transcriptional activator heat shock factor (HSF)^38, 39^. Upon heat stimulation, HSFs undergo trimerization and nuclear localization to bind to the heat shock elements (HSEs) located in the promoter region of the heat shock protein (HSP) gene, leading to the expression of HSP. We therefore utilized the HSP promoter (Hsp) in our genetic circuits to design inducible CRISPR systems.

### Inducible transgene expression regulated by heat-sensitive promoters

We tested the Hsp (HSPA7 promoter) that we and others have previously used^33, 37^, and our recently published synthetic heat-sensitive promoter 7H-YB composed of seven HSEs and a synthetic core promoter YB-TATA, which is more specific to heat stimulation^40^. In cells engineered with Hsp- or 7H-YB-driven eGFP, both heat-sensitive promoters demonstrated strong heat-inducibility, activating eGFP expression in 73.2% and 75.5% of the engineered cells with 10 min heat shock, and 92.3% and 96.4% with 20 min (HS, using a thermal cycler; Methods), respectively (Supplementary Fig. 1a-c). 7H-YB induced a mean fluorescence intensity (MFI) of eGFP approximately twice as high as that induced by Hsp, but it also caused a higher basal leakage than Hsp (11.1% vs. 0.7%, Supplementary Fig. 1c,d). Both heat-sensitive promoters were used throughout our designs and specified in the corresponding plasmid schematics. All HS experiments in this study were performed at 43 °C unless otherwise specified.

### Activation of heat-inducible genes using an in-house built FUS system

We developed an in-house built FUS system with real-time feedback temperature control for generating localized hyperthermia in vitro on cells as well as in vivo in mice (Supplementary Fig. 2a-e, Methods). A thermocouple was used to measure the FUS focal temperature, providing feedback for a PID controller to maintain the focal temperature at the target value by adjusting the FUS power (Supplementary Fig. 2a). Stable heating at 43 °C was achieved using this FUS system (Supplementary Fig. 2f). We also generated subcutaneous tumours engineered with Hsp-driven Fluc in mice and applied FUS stimulation on the tumours. Luminescence was quantified before and 6 h after FUS via IVIS, and the ratio of the after/before readings was used to indicate the induction level. We observed a 11.2-fold induction in mice with FUS and minimal induction (1.2-fold) in the ones without FUS (Supplementary Fig. 2g,h), demonstrating the capability of the in-house built FUS system in activating heat-inducible genes.

### Inducible upregulation of exogenous and endogenous genes via FUS-CRISPRa

To engineer a FUS-CRISPRa system, we adopted the Ribozyme-gRNA-Ribozyme (RGR) strategy utilizing self-cleaving HH and HDV ribozymes that enables gRNA production from inducible RNA polymerase II promoters like Hsp^41, 42^. Upon FUS/heat stimulation, Hsp initiates production of the HHRibo-sgRNA-HDVRibo transcript, which undergoes self-cleavage to generate the sgRNA (Fig. 1a). The sgRNA then integrates with the constitutively expressed dCas9 and transcriptional factors (e.g., VP64, SAM^43^) to activate target gene expression (Fig. 1a).

**Fig. 1.**
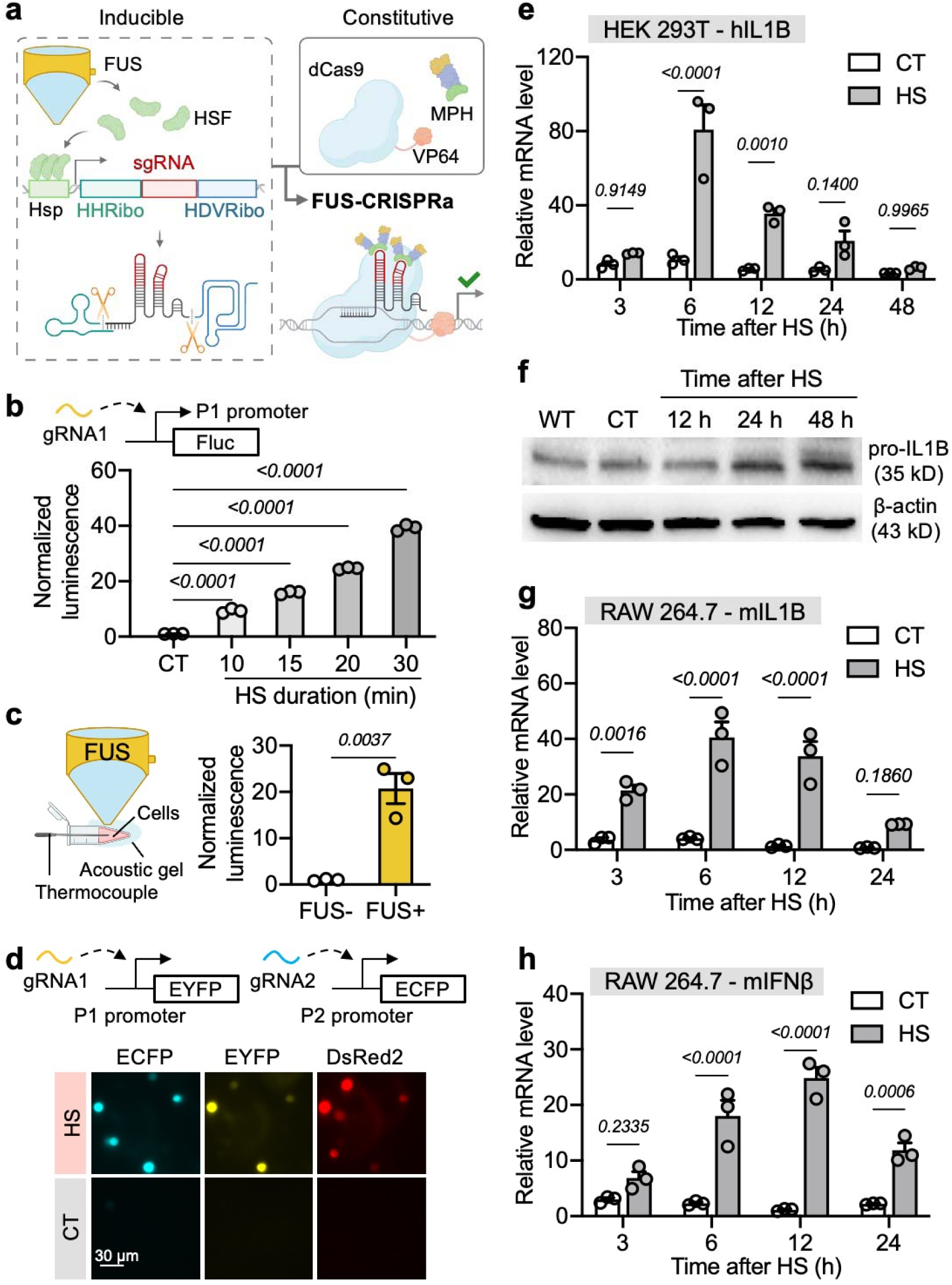
FUS-CRISPRa enables inducible upregulation of exogenous and endogenous genes. **a**, Schematic illustration of the FUS-CRISPRa system. **b**, Normalized Fluc luminescence in cells engineered with P1-targeting FUS-CRISPRa and P1-driven Fluc quantified 24 h after different durations of HS. Readings were normalized to the CT group. **c**, Left, schematic illustration of FUS stimulation of cells in vitro; Right, normalized Fluc luminescence in cells engineered with P1-targeting FUS-CRISPRa and P1-driven Fluc quantified 24 h after FUS. Readings were normalized to the FUS-group. **d**, Cells engineered with P1- and P2-targeting FUS-CRISPRa, P1-EYFP, and P2-ECFP were imaged 24 h after HS. Scale bar = 30 μm. **e**, Relative IL1B mRNA expression in HEK 293T cells engineered with hIL1B-targeting FUS-CRISPRa, normalized to IL1B mRNA level in wild type (WT) HEK 293T cells. **f**, Pro-IL1B protein expression in wild type (WT) cells or engineered cells in **e**. **g**,**h**, Relative IL1B (**g**) or IFNβ (**h**) mRNA expression in RAW 264.7 cells engineered with FUS-CRISPRa targeting mouse IL1B (**g**) or IFNβ (**h**) gene, normalized to the corresponding mRNA levels in WT RAW 264.7 cells. In **b**, CT, control, without HS; data are technical triplicates representative of three independent experiments. In **c**, FUS+, with 20 min FUS stimulation at 43 °C; FUS-, without FUS stimulation; n = 3 biological replicates. In **d**-**h**, HS, with 30 min HS; CT, without HS. In **e**, **g**, and **h**, bar heights represent means; error bars represent s.e.m.; n = 3 technical replicates representative of two individual experiments. Unpaired t test was used in **c**, two-way ANOVA followed by Sidak’s multiple comparisons test was used in **b**, **e**, **g**, **h**.

We first tested the capability of FUS-CRISPRa in activating exogenous genes. In cells transfected with FUS-CRISPRa for the inducible expression of gRNA1 targeting a synthetic promoter P1^42^ (Supplementary Fig. 3a), different durations of HS induced tunable expression of P1-driven firefly luciferase (Fluc, Fig. 1b). FUS stimulation (43 °C, 20 min) also induced a comparable level of Fluc aviation in the engineered cells in vitro (Fig. 1c). We further applied FUS in vivo in mice and observed significant Fluc activation via FUS-CRISPRa as well (Supplementary Fig. 3b). In addition, we engineered cells with multiplexed FUS-CRISPRa containing Hsp-DsRed2-RG1R-RG2R, allowing simultaneous inducible production of gRNA1 and gRNA2 targeting synthetic promoters P1 and P2 respectively^42^ (Supplementary Fig. 3c). Along with the Hsp-driven DsRed2 expression, the activations of P1-driven EYFP and P2-driven ECFP via FUS-CRISPRa were also observed in the cells with HS, with minimal background signals in control (CT) cells without HS (Fig. 1d). These results validated the design of FUS-CRISPRa with inducible gRNAs.

We then applied FUS-CRISPRa to target the genome to regulate endogenous gene expressions. We constructed an all-in-one piggyBac plasmid containing Hsp-RGR targeting the human IL1B (hIL1B) gene, which is a common target of CRISPRa^44^, together with the constitutive dCas9-SAM (Supplementary Fig. 3d) to generate cell lines accordingly (Methods). Quantification of hIL1B mRNA level and pro-IL1B protein expression in the engineered HEK 293T cells at different time points after HS revealed a trend of heat-inducible upregulation of hIL1B through FUS-CRISPRa (Fig. 1e,f). No heat-inducibility of hIL1B was observed in wild type (WT) cells (Supplementary Fig. 3e). We also validated our design in mouse RAW 264.7 cells using FUS-CRISPRa targeting mouse IL1B (mIL1B) and IFNβ (mIFNβ) genes (Fig. 1g,h). Heat itself did not significantly alter mIL1B and mIFNβ expression in WT RAW 264.7 cells (Supplementary Fig. 3f,g). In summary, FUS-CRISPRa allows inducible activation of exogenous and endogenous genes in different cell types.

We also designed a different FUS-CRISPRa system with an inducible dCas9 incorporating the SunTag system^45^. This FUS-CRISPRa system is composed of an inducible dCas9 fused to eight repeats of GCN4, a constitutive αGCN4-scFv-fused VP64^46^, and a constitutive gRNA (Supplementary Fig. 4a,b). We tested this design in activating the P1-driven Fluc (Supplementary Fig. 4c). HS robustly induced 4-6-fold of Fluc activation (HS vs. CT) in multiple cell types (Supplementary Fig. 4d). FUS also induced a comparable level of activation in vitro (Supplementary Fig. 4d) and in vivo (Supplementary Fig. 4e), validating this design of FUS-CRISPRa.

### FUS-CRISPRi-mediated epigenetic regulation for gene repression

We next sought to engineer FUS-CRISPRi for controllable gene repression for lasting periods through epigenetic reprogramming. CIRSPRoff is an epigenetic memory writer composed of dCas9, DNA methyltransferase DNMT3A-3L domains, and KRAB domains reported to durably silence gene expression^47^ (Supplementary Fig. 5a). We co-transfected HEK 293T cells with CRISPRoff and Hsp-RGR containing gRNA targeting ARPC2, a common target of CRISPRi^48^, to test heat-inducible gene repression. However, we did not observe significant ARPC2 downregulation after HS (Supplementary Fig. 5b). We also tested Hsp-RGR containing Zap70-targeting gRNA in Jurkat cells by electroporation, yet still did not observe Zap70 downregulation (Supplementary Fig. 5b). On the contrary, robust gene repression was observed when constitutive ARPC2 or Zap70 gRNA was co-transfected with CRISPRoff (Supplementary Fig. 5c). We surmised that the copy number of gRNA generated from Hsp-RGR after HS was not sufficient to induce gene repression with CRISPRoff.

Therefore, we employed a different strategy to engineer FUS-CRISPRi by changing the inducible component from gRNA to dCas9 while incorporating the SunTag amplification system as described above^44^. Since heat-inducible expression may result in a lower protein copy number than constitutive expression, we reasoned that having a heat-inducible dCas9-nxGCN4 and a constitutive scFv-regulator would allow a favorable stoichiometry to promote the recruitment of multiple copies of the regulators to a given dCas9 complex. As such, this FUS-CRISPRi system is composed of the 7H-YB promoter (stronger induction capability than the Hsp, Supplementary Fig. 1) driving the dCas9 fused to eight repeats of GCN4, a constitutive EFS promoter driving a previously reported αGCN4-scFv-fused epigenetic regulator DNMT3A-3L, and the constitutive U6 promoter driving the gRNA^49^ (Fig. 2a, Supplementary Fig. 6a). FUS stimulation can induce dCas9-8xGCN4 expression, allowing the recruitment of multiple copies of the epigenetic regulators through the scFv. As such, the complex is brought to the target locus by the gRNA to repress gene expression via DNA methylation (Fig. 2a).

**Fig. 2.**
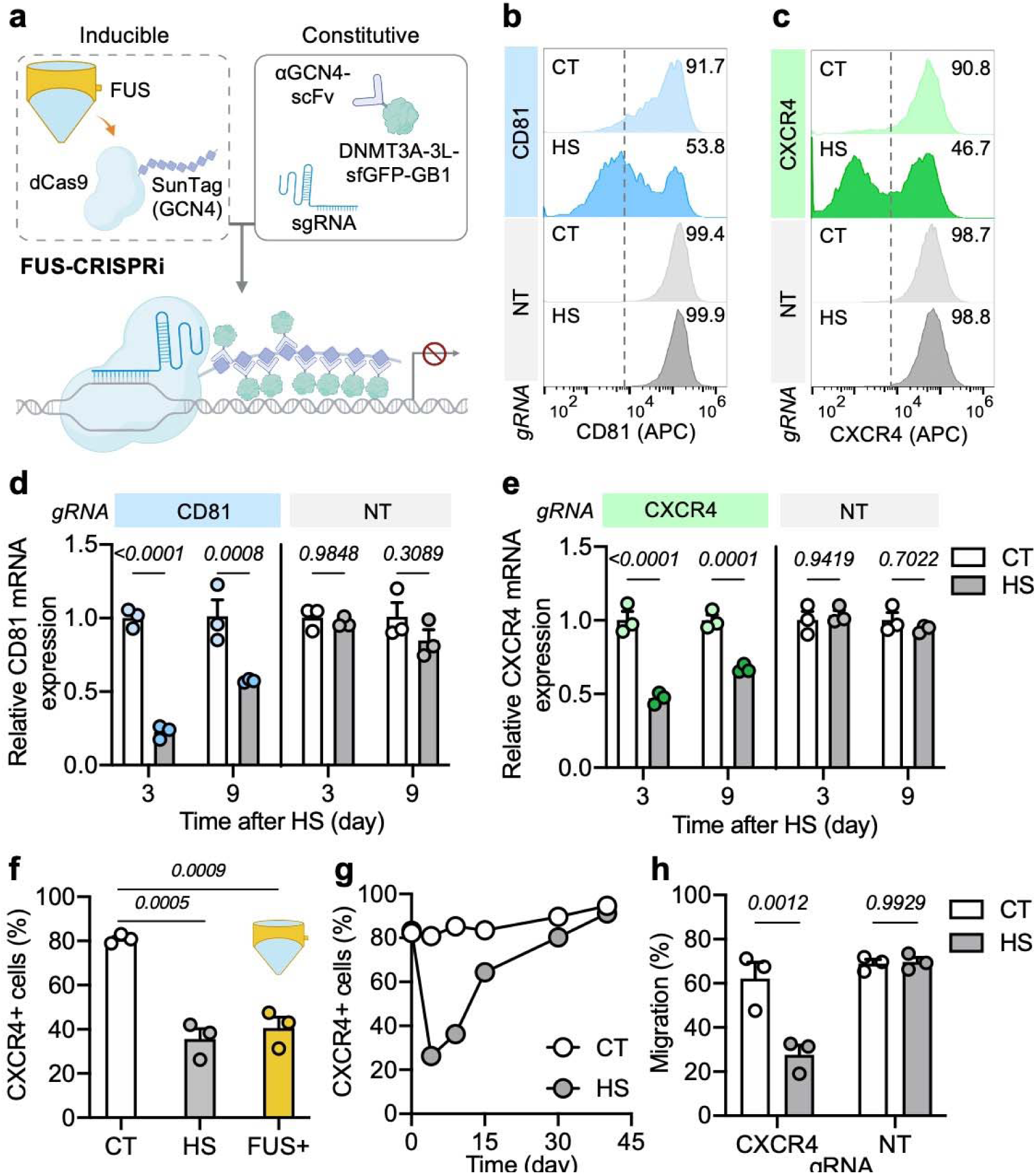
FUS-CRISPRi-mediated inducible suppression of endogenous genes. **a**, Schematic illustration of the FUS-CRISPRi system. **b**,**c**, Representative flow cytometry data of CD81 (**b**) or CXCR4 (**c**) expression in FUS-CRISPRi-engineered Jurkat cells with gRNA targeting CD81 (**b**) or CXCR4 (**c**), or with non-targeting (NT) gRNA. The cells were stained with anti-CD81 (**b**) or anti-CXCR4 (**c**) antibody four days after HS. **d**, Relative CD81 mRNA expression 3 or 9 days after HS in cells in **b**. **e**, Relative CXCR4 mRNA expression in cells in **c**. **f**, Percentage of CXCR4+ cells in Nalm6 cells engineered with CXCR4-targeting or NT FUS-CRISPRi with DNMT mutant with different treatments. **g**, Kinetics of CXCR4 expression in cells engineered with CXCR4-targeting FUS-CRISPRi. **h**, The migration ability (%) of the engineered FUS-CRISPRi Nalm6 cells in a transwell assay. In **b**-**h**, HS, with 20 min HS; CT, without HS. In **f**, FUS+; with 20 min FUS stimulation at 43 °C on cells in vitro. In **d** and **e**, bar heights represent means of technical triplicates representative of two individual experiments. In **h**, bar heights represent means of biological triplicates. Error bars represent s.e.m. Two-way ANOVA followed by Sidak’s multiple comparisons test was used for statistical analysis.

We transduced Jurkat cells with the FUS-CRISPRi system containing gRNAs targeting surface markers CD81 or CXCR4, which can be quantified by staining. Cell surface staining of CD81 four days after HS showed a significant decrease in CD81 expression in the HS cells compared with non-heated control (CT) cells (53.8% vs. 91.7%, Fig. 2b). Similarly, CXCR4 expression was also repressed by HS (46.7% in HS vs. 90.8% in CT cells, Fig. 2c). HS itself did not affect CD81 or CXCR4 expression in the cells with non-targeting NT gRNA (Fig. 2b,c). The effect of FUS-CRISPRi-mediated gene repression was also confirmed by quantification of the corresponding mRNA levels (Fig. 2d,e). Similar gene repression effects were achieved in Nalm6 cells engineered with FUS-CRISPRi (Supplementary Fig. 6b-d).

CXCR4 is a chemokine receptor known to promote tumour growth and metastasis^50–52^. We therefore examined the effect of FUS-CRISPRi-mediated CXCR4 downregulation in Nalm6 tumour cells. We also replaced the WT DNMT in the original FUS-CRISPRi with a previously reported DNMT mutant of reduced off-target methylation (Supplementary Fig. 7a). A dramatic reduction of CXCR4 expression was seen in CXCR4 FUS-CRISPRi cells four days after HS compared with those without HS, and FUS stimulation was able to induce a comparable repression effect in the engineered cells (Fig. 2f). Dynamic tracking revealed that the CXCR4 expression in the cells with HS recovered to a level similar to that in the cells without HS in approximately 40 days, indicating a sustained but reversible effect of FUS-CRISPRi (Fig. 2g, Supplementary Fig. 7b). Transwell assays further demonstrated that the migration ability was compromised in cells with HS-induced CXCR4 downregulation (Fig. 2h). Taken together, our results suggest that FUS-CRISPRi allows inducible and reversible gene repression on different genes through epigenetic modulation in different cell types, allowing the control of cellular functions by ultrasound.

### FUS-CRISPR-mediated knockout of endogenous genes

One of the advantages of the FUS-inducible system is its ability to transiently activate regulators (e.g., Cas9) that may be immunogenic or toxic if expressed constitutively^12^. Following the development of FUS-CRISPRa and FUS-CRISPRi, we engineered FUS-CRISPR composed of inducible Cas9 and constitutive gRNAs (Fig. 3a, Supplementary Fig. 8a,b) and verified heat-inducible Cas9 expression in the engineered cells (Fig. 3b). In Jurkat T cells engineered with FUS-CRISPR targeting key signaling molecules CD3D or Zap70, HS induced CD3D knockout (KO) in 44.3% cells and Zap70 KO in 39.2% cells as quantified by genotyping PCR and sequencing (Fig. 3c). Low levels of basal KO were observed in CT cells (13% for CD3D and 15.4% for Zap70), likely due to the leakage of the heat-sensitive promoters (Fig. 3c). To test whether HS-induced KO can affect cellular functions, we stimulated the Jurkat T cells with anti-T-cell receptor (TCR) antibody and quantified T-cell activation by CD69 staining. As expected, since CD3D is a subunit of the TCR complex and Zap70 is a critical mediator of the TCR signaling pathway, Jurkat cells with HS-induced KO of CD3D or Zap70 demonstrated significantly weakened TCR-dependent T-cell activation, reflected by CD69 expressions (Fig. 3d).

**Fig. 3.**
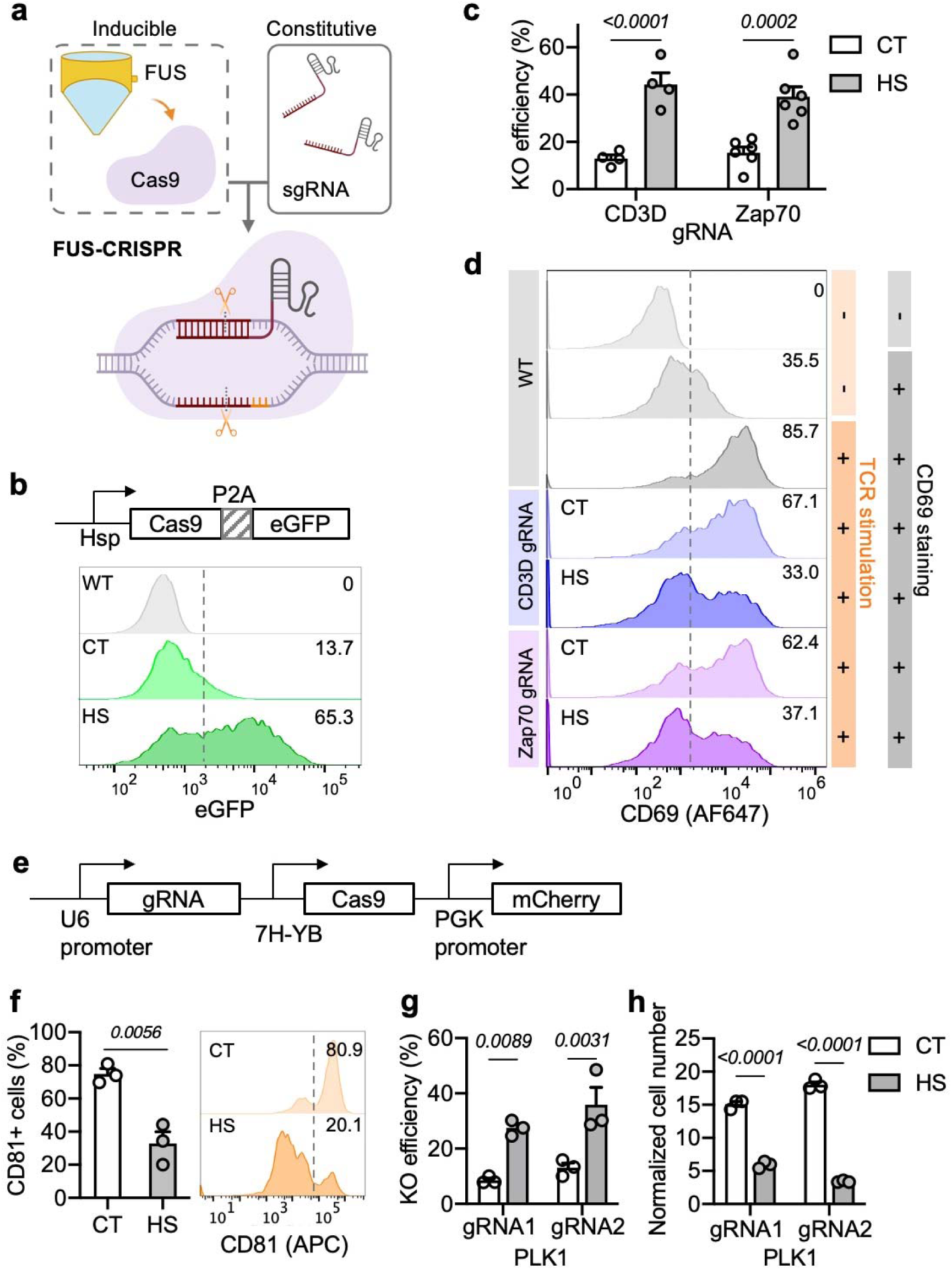
FUS-CRISPR-mediated knockout of target genes. **a**, Schematic illustration of the FUS-CRISPR system. **b**, Heat-inducible Cas9 expression represented by eGFP signal under flow cytometry in engineered Jurkat cells. **c**, Knockout efficiencies in Jurkat cells engineered with FUS-CRISPR targeting CD3D or Zap70 quantified four days after HS. N = 4 and 6 biological replicates for CD3D and Zap70, respectively. **d**, CD69 staining of WT or FUS-CRISPR-engineered Jurkat cells after TCR stimulation. **e**, The all-in-one FUS-CRISPR plasmid. **f**, Percentage of CD81+ cells (left) and the representative flow cytometry profile (right) in U-87 MG cells engineered with CD81-targeting FUS-CRISPR quantified 8 days after HS. **g**, Knockout efficiencies in Nalm6 cells engineered with FUS-CRISPR with different gRNAs targeting PLK1 gene, quantified four days after HS. **h**, Normalized cell number of the cells in **g** on Day 4 after HS. Cell number was normalized to Day 0. In **c**, **d**, and **f**, HS, with 20 min HS; CT, without HS. In **g** and **h**, HS, with 15 min HS; CT, without HS. Bar heights represent means; error bars represent s.e.m. In **f** and **g**, n = 3 biological replicates. In **h**, n = 3 technical replicates representative of two independent experiments. Unpaired t test was used in **f**, two-way ANOVA followed by Sidak’s multiple comparisons test was used in **c**, **g**, **h**.

To examine the feasibility of broad applications, we further engineered an all-in-one plasmid for FUS-CRISPR and tested it in multiple tumour cell lines (Fig. 3e). Surface staining of U-87 MG glioma tumour cells engineered with CD81-targeting FUS-CRISPR showed that HS induced significant CD81 KO (Fig. 3f). To explore the therapeutic applications of FUS-CRISPR, we generated Nalm6 tumour cells containing FUS-CRISPR targeting polo-like kinase 1 (PLK1, Supplementary Fig. 8c), a key regulator of cell cycle and an active target of cancer therapy^18, 53^. HS induced PLK1 KO and significantly inhibited cell proliferation with different PLK1-targeting gRNAs (Fig. 3g,h). In summary, FUS-CRISPR can be applied to control genome editing of endogenous genes and reprogramming of cellular functions.

### Telomere disruption by FUS-CRISPR

In addition to genetic editing of single genes, we hypothesized that FUS-CRISPR can act with a higher editing efficiency on repetitive loci such as telomeres than on non-repetitive loci. It has been reported that telomere dysfunction can trigger catastrophic events leading to cell senescence and apoptosis^54–56^. We hence co-transfected HEK 293T cells with FUS-CRISPR containing the gRNA targeting repetitive telomere sequences (Supplementary Fig. 8d) and HaloTag-fused 53BP1, a marker for DNA double strand breakage (DSB) to report the genome editing sites. Fluorescence microscopy revealed that HS induced DSB at multiple loci in the cells with telomere-targeting FUS-CRISPR, as evidenced by the dotted 53BP1 pattern, which was not observed in non-activated CT cells or cells with non-targeting NT FUS-CRISPR (Fig. 4a). We also co-transfected the cells with tagBFP-fused telomeric repeat binding factor 2 (TRF2) to mark the telomere loci^57^. Merged images of 53BP1 and TRF2 showed multiple colocalization puncta, confirming the presence and precision of FUS-CRISPR-induced DSB at telomeres (Fig. 4a).

**Fig. 4.**
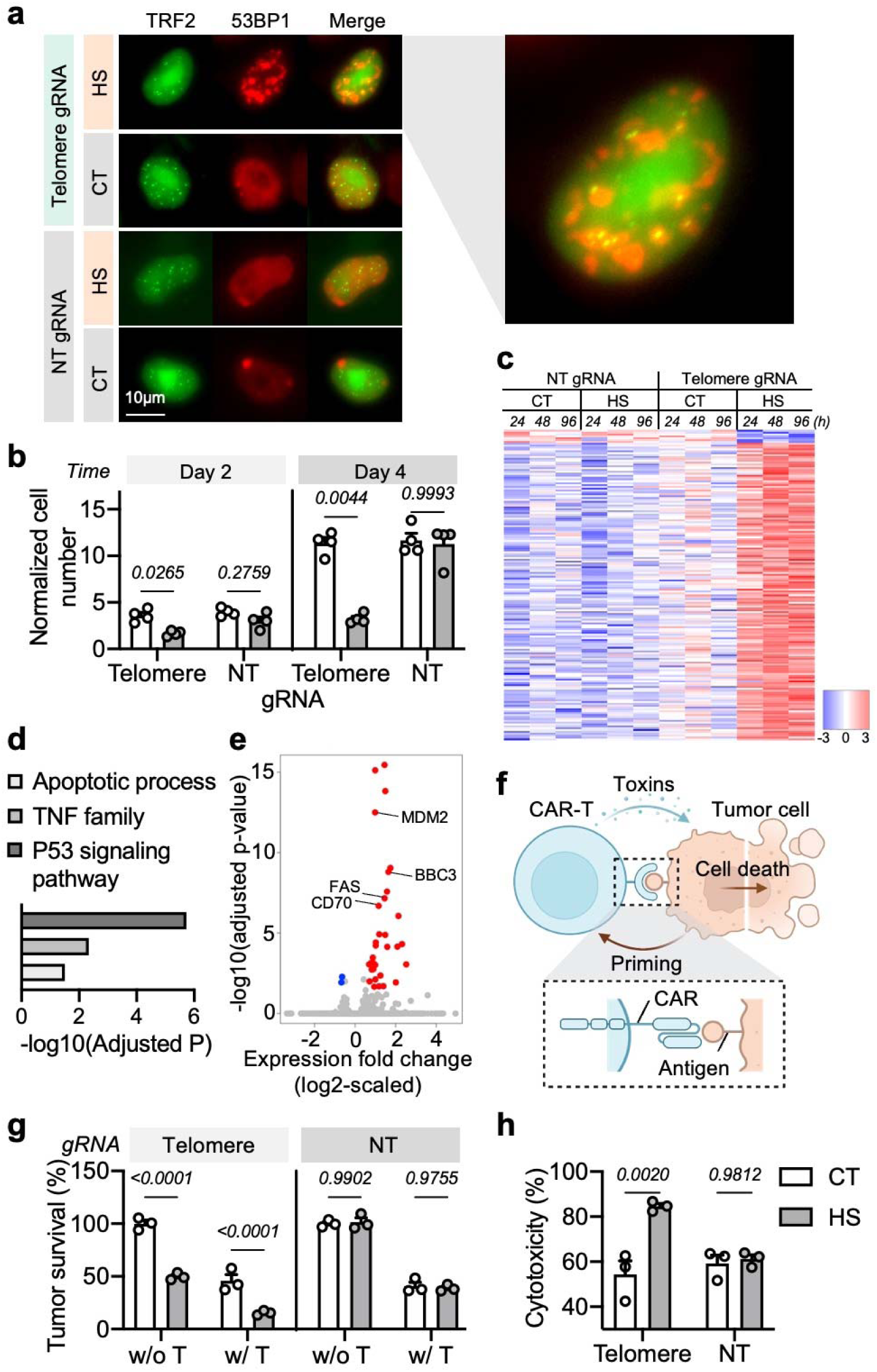
FUS-CRISPR-mediated telomere disruption can inhibit tumour cell growth and its resistance to CAR-T cell killing. **a**, Nuclear distribution of tagBFP-TRF2 and HaloTag-53BP1 in FUS-CRISPR-engineered HEK 293T cells with telomere-targeting gRNA or non-targeting (NT) gRNA. HS, with 30 min HS; CT, without HS. Right, enlarged image merging TRF2 and 53BP1 signals. Scale bar = 10 μm. **b**, Normalized cell number of FUS-CRISPR-engineered Nalm6 cells with telomere-targeting gRNA or NT gRNA two (D2) or four (D4) days after HS. Cell number was normalized to Day 0. N = 4 biological replicates. **c**, Heat-map of differential gene expression in Nalm6 cells engineered with telomere-targeting or NT FUS-CRISPR at 24, 48, or 96 h after HS. **d**, The top three enriched GO terms in the HS group compared to the CT group in the telomere-targeting FUS-CRISPR cells in **c**. **e**, Volcano plot showing the downregulated (blue) and upregulated (red) genes between HS and CT groups in the telomere-targeting FUS-CRISPR cells in **c**. **f**, Schematic illustration of CAR-T cell attack on tumour cells. **g**, Survival (%) of FUS-CRISPR-engineered Nalm6 tumour cells 72 h after culture with (w/T) or without (w/o T) αCD19CAR-T cells in the luciferase-based cytotoxicity assay. The survival (%) was normalized to CT, w/o T group. **h**, Cytotoxicity (%) of CAR-T cells in the co-culture groups (w/ T) in **g**. The cytotoxicity (%) was quantified as 100%_j–_jTumour survival (%). In **g** and **h**, n = 3 technical replicates. Data are representative of two independent experiments. In **b**, **c**, **g**, and **h**, HS: with 10 min HS; CT, without HS. Bar heights represent means; error bars represent s.e.m. Two-way ANOVA followed by Sidak’s multiple comparisons test.

We then engineered Nalm6 tumour cells with telomere-targeting or NT FUS-CRISPR. Consistent with previous reports of telomere-dysfunction-related cell senescence and apoptosis, we observed that a relatively short duration of HS (10 min) significantly inhibited the proliferation of the cells engineered with telomere FUS-CRISPR, but not that of the cells with NT FUS-CRISPR, suggesting that telomere disruption rather than hyperthermia itself suppressed cell growth (Fig. 4b). Bulk RNA-seq further revealed that FUS-CRISPR-mediated telomere disruption led to the upregulation of multiple genes associated with the stress response p53 signaling pathway and apoptotic process (e.g., MDM2, FAS, BBC3) and the TNF family (e.g., CD70) in the engineered cells to trigger cell cycle arrest (Fig. 4c-e, Supplementary Fig. 9)^58^. This priming effect of FUS-CRISPR on tumour cells may hence not only cause the tumour cell cycle arrest and apoptosis, but also induce T cell immune responses via TNF family^59^.

To test whether telomere disruption affect tumour killing by T cells, we employed anti-CD19 chimeric receptor antigen (CAR)-T cells specifically targeting CD19^+^ Nalm6 tumour cells (Fig. 4f, Supplementary Fig. 10). Fluc-expressing FUS-CRISPR Nalm6 cells with or without HS were co-cultured with CAR-T cells at a low effector-to-target (E:T) ratio of 1:20 for luciferase-based killing assay. The percentage of surviving tumour cells and the corresponding cytotoxicity of the CAR-T cells were quantified from Fluc luminescence 72 h after co-culture (Fig. 4g,h). CAR-T cells demonstrated significantly stronger cytotoxicity against Nalm6 cells with HS-induced telomere disruption than that against CT Nalm6 cells (84.6% vs. 54.3%), while similar cytotoxicities were observed against NT FUS-CRISPR Nalm6 cells with or without HS (59.2% and 61.2%, respectively, Fig. 4h). These results indicated that tumour cells with induced priming and telomeric DSB were less resistant to CAR-T cell killing.

### FUS-CRISPR-mediated telomere disruption aids CAR-T therapy for tumour treatment

Encouraged by the effect of FUS-CRISPR-mediated telomere disruption in vitro, we investigated its therapeutic potentials in vivo. We generated subcutaneous tumours in NSG mice using Fluc^+^ Nalm6 cells engineered with telomere FUS-CRISPR or NT FUS-CRISPR. The tumours were treated with (FUS+) or without (FUS-) 10 min FUS on Days 9 and 12 (Fig. 5a and Supplementary Fig. 11a). No significant difference in growth was observed between NT FUS-CRISPR tumours with or without FUS, indicating that FUS alone did not affect tumour growth (Supplementary Fig. 11b-e). In the mice bearing telomere FUS-CRISPR tumours, FUS+ tumours exhibited mildly inhibited growth compared with the FUS-tumours from bioluminescence imaging (BLI) yet no statistically significant difference from caliper measurement (Fig. 5b-d). Both the FUS+ and FUS-groups showed 0% survival at the end of observation (Fig. 5e). These results suggested that FUS-CRISPR-mediated telomere disruption alone was not sufficient for tumour treatment.

**Fig. 5.**
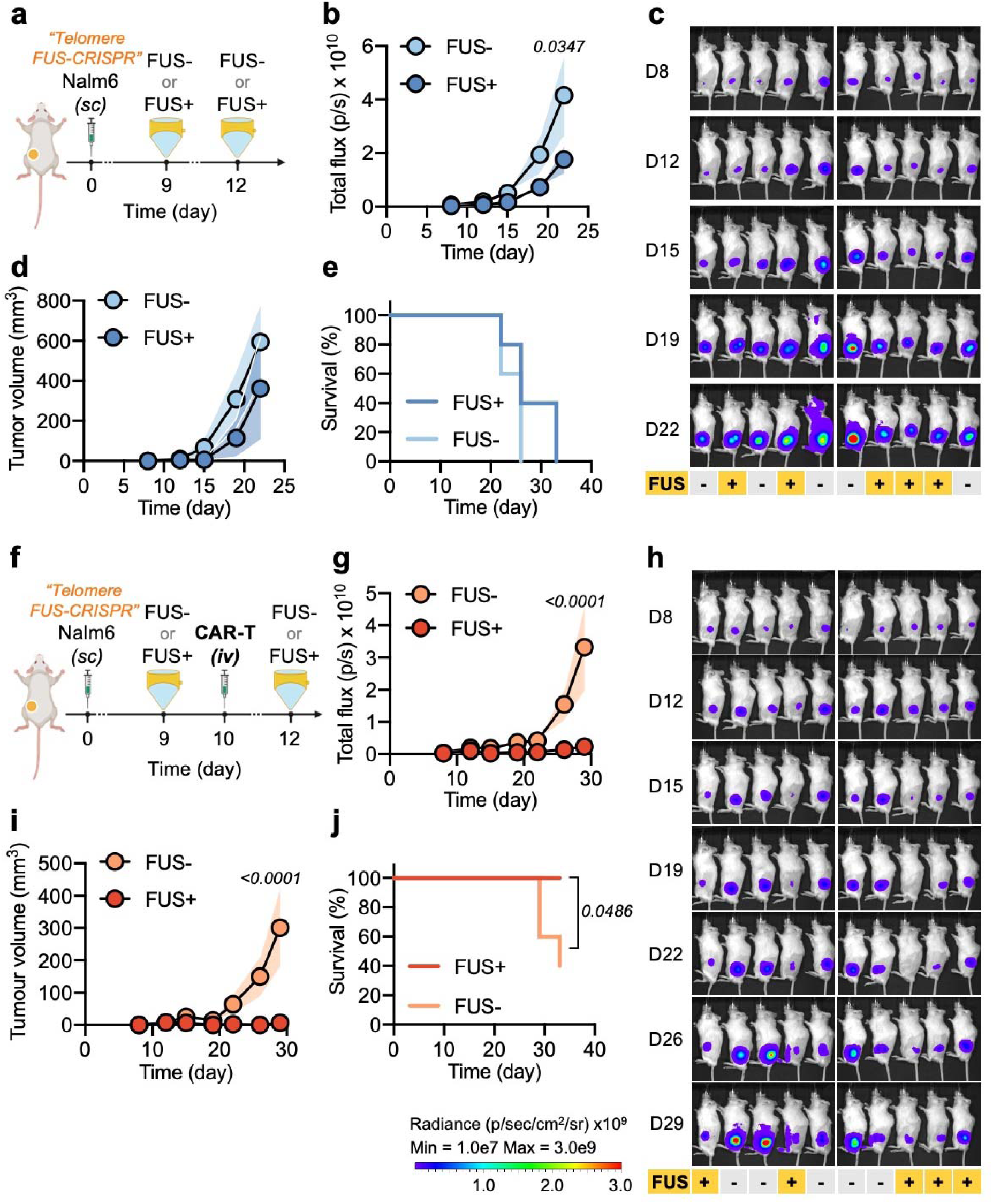
FUS-CRISPR-mediated telomere disruption enhances the efficacy of CAR-T therapy in vivo. **a**, Timeline of FUS-CRISPR-mediated telomere disruption experiment in NSG mice. **b-d**, Tumour aggressiveness in the mice in **a** quantified by total flux of the tumour from BLI measurement (**b**), the corresponding BLI images (**c**), and the tumour volume based on caliper measurement (**d**). **e**, Survival curves of the tumour-bearing mice in **a**. **f**, Experimental timeline of FUS-CRISPR combined with CAR-T therapy in NSG mice. **g-i**, Tumour aggressiveness in the mice in **f** quantified by total flux of the tumour (**g**), the corresponding BLI images (**h**), and the caliper-measured tumour volume (**i**). **j**, Survival curves of the tumour-bearing mice in **f**. Data points represent means; error bands represent s.e.m.; n = 5 mice per group. Two-way ANOVA followed by Sidak’s multiple comparisons test was used in **b**, **d**, **g**, and **i**. Log-rank (Mantel-Cox) test was used in **e** and **j**.

Therefore, we hypothesized that a treatment strategy combining FUS-CRISPR-mediated telomere disruption for tumour priming and CAR-T therapy could synergistically lead to a more prominent therapeutic outcome. We accordingly generated subcutaneous tumours in mice using telomere FUS-CRISPR Nalm6 cells followed with (FUS+) or without (FUS-) FUS stimulation (Fig. 5f). Ten days later, we injected a low dose of CAR-T cells intravenously in both FUS+ and FUS-groups (Fig. 5f). We observed significantly suppressed growth of the tumours in the FUS+ group compared to that of FUS- (Fig. 5g-i). The two groups of mice also showed different survival profiles: while all the mice in the FUS+ group survived, only 40% (two out of five) mice in the FUS-group responded to CAR-T therapy, and the rest 60% mice had reached euthanasia criteria due to tumour progression by the end of observation (Fig. 5j). We further performed a control experiment using NT FUS-CRISPR tumours with CAR-T treatment in both FUS- and FUS+ groups (Supplementary Fig. 11f). There was only a mild inhibition of tumour growth in the FUS+ group compared with the FUS-group, but there was no significant difference in the survival rate between the two groups (Supplementary Fig.11g-j). Taken together, telomere-targeting FUS-CRISPR can allow ultrasound-controllable genome editing and tumour priming for efficient CAR-T therapy to achieve synergistic therapeutic effects.

## Discussion

We developed a FUS-CRISPR(a/i) toolbox including FUS-controllable CRISPRa, CRISPRi, and CRISPR systems that allowed inducible control of genetic and epigenetic reprogramming by FUS. We demonstrated inducible upregulation, downregulation, and knockout of exogenous and/or endogenous genes in multiple cell types in vitro and in vivo using FUS. We further induced multiple DSBs at telomere sites in tumour cells via telomere-targeting FUS-CRISPR, which primed tumours for efficient killing by cytotoxic CAR-T cells in vitro and in vivo. The combined strategy demonstrated synergistic therapeutic effects and may promote CAR-T therapy against relatively resistant tumours via priming.

Ultrasound and its integration with genetic engineering and synthetic biology have revolutionized the control of genetics and cellular functions in live animals with unprecedented penetration depth at tens of centimeters^37, 60, 61^. Despite its high temporal resolution (e.g., hundreds of frames per second), the spatial resolution of traditional ultrasound is however limited at submillimeter levels^62^. With recent development in acoustic reporter genes (ARGs) and functional ultrasound localization microscopy, ultrasound imaging can achieve spatial resolutions in micrometers and at single cell levels^63–65^. Similarly, it is expected that the ultrasound control of genetics and cellular functions can reach the level of single cells and subcellular compartments. The FUS-CRISPR(a/i) toolbox developed in this work can further allow the ultrasound-guided regulation in the dimensions of genome and epigenome at single-base precision^66^. As such, the FUS-CRISPR(a/i) toolbox should provide a versatile platform to allow the remote and noninvasive control of genome and epigenome in specific tissues/organs of genetically engineered animals with high spatiotemporal resolution.

Adeno-associated virus (AAV) has been demonstrated to allow gene delivery in animals and humans with high efficiencies^67, 68^. We envision that, in the future, the FUS-CRISPR(a/i) cassettes can be directly delivered in vivo using AAV followed by FUS-induced localized hyperthermia to activate CRISPR(a/i) in living organisms. Transgenic FUS-CRISPR(a/i) mouse models similar to tet-controllable Cas9 mice^69, 70^ may also be developed. Such advancements should fully unleash the power of FUS-controllable technologies for genomic manipulation in live animals and patients in a remote, noninvasive, and spatiotemporally precise fashion. The FUS-CRISPR(a/i) technology should benefit fundamental, translational, and clinical research, with its applications ranging from interrogation of gene functions in targeted tissues/locations and/or CRISPR screening under physiological context in transgenic mice, to disease treatment in specific tissues in patients.

CRISPR-Cas9 proteins have been a powerful tool for genome editing, but can evoke adaptive immune responses and tissue damages in vivo, and are therefore potentially pathogenic if used to correct inherited genetic defects to treat diseases^71^. Protein engineering to remove immunogenic epitopes and humanize these synthetic proteins to circumvent this issue can be difficult owing to the high diversity of the human leukocyte antigen (HLA) loci^72^. Using our acoustogenetics approach, the transiently induced genomic and epigenomic regulators can be cleared in a timely manner to mitigate or evade the adaptive immune response, offering a new option for genome editing and gene therapy at specific tissues/organs. Indeed, FUS-CRISPR-mediated telomere editing allowed tumour priming via upregulations of multiple genes associated with tumour cell apoptosis and immune activation, which enhanced the efficacy of CAR-T therapy. We reasoned that the targeting of repetitive telomeric sequences may lead to a higher editing efficiency than targeting a single gene. While we tested tumour priming in Nalm6 lymphoma cells in this work, the modular design of FUS-CRISPR(a/i) should allow the general genome/epigenome regulation and priming in other types of tumour cells. We anticipate that this tumour priming strategy can be broadly applied to aid CAR-T therapy against more resistant tumour types^73–75^. FUS can control the reprogramming to occur only at tumour regions for precise and safe tumour eradication.

In summary, the FUS-CRISPR(a/i) toolbox developed here adds to the collection of FUS-based acoustogenetics technologies. FUS-CRISPR(a/i) can be integrated with different CRISPR regulators and gRNAs, and such a modular design should enable targeting of, in principle, any accessible genomic locus for various reprogramming purposes. FUS-CRISPR(a/i) can also be used for tumour priming and synergistically combined with other therapies such as CAR-T therapy for effective cancer treatments.

## Methods

### General cloning

Plasmids were constructed by Gibson Assembly (NEB, E2611L), T4 ligation (NEB, M0202L), or Golden Gate Assembly. PCR was performed using synthesized primers (Integrated DNA Technologies) and Q5 DNA polymerase (NEB, M0491). The sequences of the constructed plasmids were verified by Sanger sequencing (Azenta). Plasmids used in this study and their corresponding templates are listed in Supplementary Table 1. The sequences of the gRNAs were obtained from literature and listed in Supplementary Table 2^43, 44, 47, 48, 55, 76–78^.

### General cell culture and antibodies

HEK 293T and RAW 264.7 cells were cultured in Dulbecco’s Modified Eagle’s Medium (DMEM) (Gibco, 10569010) supplemented with 10% fetal bovine serum (FBS) (Gibco, 10438026) and 1% penicillin– streptomycin (P/S) (Gibco, 15140122). Jurkat and Nalm6 cells were cultured in Roswell Park Memorial Institute Medium (RPMI 1640) (Gibco, 22400105) with 10% FBS and 1% P/S. Primary human T cells were cultured in complete RPMI 1640 supplemented with 100□U/ml recombinant human IL-2 (PeproTech, 200-02). All mammalian cells were cultured at 37□°C in a humidified 5% CO_2_ incubator.

The antibodies used in this study are listed in Supplementary Table 3.

### Gene delivery methods

General plasmid transfection in HEK 293T cells were performed using Lipofectamine 3000 transfection reagent (Invitrogen, L3000001) according to the manufacturer’s protocol.

Electroporation in Jurkat cells was performed as previously described^79^. Briefly, ten million Jurkat cells were resuspended in 500 μl of OptiMEM containing 20 μg Hsp-RGR or U6-gRNA plasmid and 20 μg CRISPRoff plasmid (Supplementary Fig. 5b-c) in a 4-mm cuvette and electroporated at 270 V, 950 μF (exponential wave, infinite resistance) using the Bio-Rad Gene Pulser Xcell Electroporation System. Cells were transferred to prewarmed culture media immediately after electroporation.

For piggyBac-based cell line generation (Fig. 1e-h), the piggyBac transposon vector (Supplementary Fig. 3d) and the piggyBac transposase plasmid (SBI, PB210PA-1) were delivered into cells at a ratio of 2.5:1 by Lipofectamine transfection in HEK 293T cells or by electroporation in Raw 264.7 cells using the Lonza 4D-Nucleofector and the SF kit (Lonza, V4XC-2032). Puromycin selection (5 μg/ml) was applied for 10 days.

For lentiviral transduction, the lentivirus was produced by transfecting HEK 293T cells with the transfer plasmid, packaging plasmid, and envelope plasmid using calcium phosphate-mediated transfection method (Promega, E1200) and harvesting the supernatant 48 - 72 h after transfection. For transduction of cell lines, 100-500 μl of unconcentrated lenvirus was added to 1×10^5^ cells. For transduction of primary human T cells, the lentivirus was concentrated using Lenti-X™ Concentrator (Takara, 631232) followed by transduction as detailed in the **Isolation, culture, and lentiviral transduction of primary human T cells** section. FACS was performed to enrich the engineered cell populations when transduction efficiency was lower than 90% for cell lines or lower than 60% for primary T human cells.

### In vitro heat shock

Cells were resuspended in regular culture media in 8-strip PCR tubes with 50□μl per tube and received heat shock (HS) in a thermal cycler (Bio-Rad, 1851148) for various durations before returning to normal culture condition. All in vitro HS experiments were performed at 43□°C.

### Activation of exogenous genes via FUS-CRISPRa

For Fig. 1b, HEK 293T cells were co-transfected with three FUS-CRISPRa plasmids (Supplementary Fig. 3a) at 1:1:1 ratio using Lipofectamine in a 12-well plate with 900 ng total DNA per well. Approximately 18 h after transfection, cells were resuspended in culture medium, equally aliquoted into PCR tubes, and subjected to different HS treatment. The content of each individual PCR tube was added to individual wells containing 150 μl prewarmed medium in a 96-well plate (Corning, 3904) and returned to normal cell culture condition.

The luminescence of each well was measured 24 h later using the Bright-Glo substrate (Promega, E2610) and a Tecan Infinite M200 Pro plate reader.

For Fig. 1d, HEK 293T cells were co-transfected with four FUS-CRISPRa plasmids (Supplementary Fig. 3c) at 1:1:1:1 ratio using Lipofectamine in a 12-well plate with 1 μg total DNA per well. HS was performed 18 - 24 h after transfection. Imaging was performed 24 h after HS as described in the **Fluorescence microscopy** section.

### Quantitative PCR

Total RNA was extracted from cells using Quick-RNA Microprep Kit (Zymo Research, R1050) and reverse transcribed to obtain cDNA using SuperScript™ IV Reverse Transcriptase (Invitrogen, 18090010). Quantitative PCR (qPCR) was performed using iTaq Universal SYBRRTM Green Supermix (Bio-Rad, 1725121) with primers listed in Supplementary Table 4.

### Western blot analysis

Cells/tumours were harvested and homogenized with RIPA buffer (Cell signaling Technology, 9806S) containing protease and phosphatase inhibitor cocktail (Merck, 04693116001 and 4906837001). The same amount of protein lysate was loaded into a pre-cast polyacrylamide SDS-PAGE gel (Bio Rad, 3450123) and ran at 30 mA for 90 min. The separated proteins were transferred onto 0.45 μm PVDF membrane (Bio Rad, 1620184) at 230 mA for 100 min. After blocking with TBS-T (Tris-buffer saline containing 0.1% Tween 20) containing 5% powdered milk for 60 min, membrane was incubated with primary antibodies against IL1B (Abcam, Ab2105) and β-actin (Santa Cruz, sc-69879) overnight at 4□°C subsequently and the corresponding HRP-conjugated secondary antibodies, followed by chemiluminescence detection using a Bio-Rad ChemiDoc XRS+ gel imager.

### Fluorescence microscopy

Microscopic images were taken with a Nikon Eclipse Ti inverted microscope with a cooled charge-coupled device (CCD) camera. For Fig. 1d and Supplementary Fig. 10b, HEK 293T or primary human T cells were dropped onto uncoated glass-bottom dishes (Cell E&G, GBD00002-200) followed immediately by imaging. For Fig. 4a, HEK 293T cells were resuspended in staining media (regular media containing Janelia Fluor® HaloTag® Ligands at 1:2000 dilution) and seeded onto fibronectin(Sigma Aldrich, F1141)-coated glass-bottom dishes. Three hours later, staining media were washed out three times and replaced with regular media. Images were taken 6 h after seeding.

### Transwell migration assay

7.5×10^4^ Fluc^+^ cells in 100 μl culture medium were seeded onto Polycarbonate Membrane Transwell inserts (Corning, 3422). 600 μl culture media containing 10 ng/ml CXCR4 ligand CXCL12 (Peprotech, 300-28A) were added to the transwell lower chambers as the chemoattractant. The cells in the inserts and the lower chambers were collected separately 3 h later followed by quantification of luminescence as described above.

Total luminescence of sample X = Luminescence of X insert + Luminescence of X lower chamber Migration (%) of sample X = (Luminescence of X lower chamber / Total luminescence of X) x 100%

### TCR stimulation in Jurkat cells

Jurkat cells were cultured in cell culture medium containing 1.7 μg/ml anti-TCR antibody (Sigma-Aldrich, 05-919) overnight followed by anti-CD69 antibody staining (Biolegend, 310910).

### Isolation, culture, and lentiviral transduction of primary human T cells

Human peripheral blood mononuclear cells (PBMCs) were isolated from buffy coats (Excellos) using lymphocyte separation medium (Corning, 25-072-CV), sorted with Pan T Cell Isolation Kit (Miltenyi, 130-096-535) to obtain primary human T cells, and activated by adding Dynabeads (Gibco, 11141D) at 1:1 bead-to-cell ratio. Two to three days later, T cells were mixed with lentivirus at multiplicity of infection (MOI) equal to 5 in Retronectin (Takara, T100B)-coated culture plates and centrifuged at 1800 g for 1 h at 32 °C for lentiviral transduction before returning to normal culture condition. Approximately one week later, T cells (with Dynabeads removed) were used for downstream applications or cryopreserved for future usage.

### Quantification of knockout (KO) efficiency

Genomic DNA was extracted from cells using Quick-DNA Miniprep Plus Kit (Zymo Research, D4068). An approximately 500bp fragment flanking the gRNA target site in the genome of engineered or WT cells was amplified by PCR with primers designed through NCBI Genome Data Viewer and Primer-BLAST (Supplementary Table 5). Sanger sequencing of the PCR products was performed to obtain trace files, which were uploaded to TIDE (TIDE created by Bas van Steensel lab, http://shinyapps.datacurators.nl/tide/) to quantify the KO efficiency.

### Quantification of cell proliferation in vitro

Cells were stained with a live/dead dye AOPI (Nexcelom, CS2-0106) and counted using an automated cell counter (Nexcelom, Cellometer K2) to determine the cell number before seeding (Day 0). The same number of cells were then seeded in a 24-well plate for different groups. Cell culture media were refreshed every two days. At the time points specified in the corresponding figure legends (Fig. 3h, Fig. 4b), cells were collected and counted again as described above to determine the number of live cells, which was then normalized to the seeding cell number on Day 0 to obtain the normalized cell number.

### Bulk RNA-seq

Nalm6 cells engineered with telomere-targeting or NT FUS-CRISPR were subjected to 10 min HS or no treatment (CT). Total RNA was collected at 24, 48, and 96 h after HS using the RNA microprep kit (Zymo Research, R1050) and sent for bulk RNA-seq (Novogene). RNA-seq data analysis was performed as previously described^80^. Briefly, raw RNA-seq reads were first preprocessed using Ktrim software (v1.4.1)^81^ to remove sequencing adaptors and low-quality cycles; PCR duplicates (i.e., reads with identical sequences) and ribosomal RNAs were then removed using in-house programs and the remaining reads were aligned to the human genome (build GRCh38/hg38) using STAR software (v2.7.9a)^82^; expression quantification were performed using featureCounts software (v2.0.3)^83^ against RefSeq gene annotation^84^; differential expression analysis were performed using DESeq2 software (v1.26.0)^85^; genes with an expression change larger than 1.5-fold and adjusted p-value smaller than 0.05 were considered as differentially expressed genes (DEGs). Functional annotation of the DEGs was performed using DAVID webserver^86^. RNA-seq results from the three time points (24, 48, and 96 h) in the same treatment group were considered as three repeats for data analysis in Fig. 4d,e and Supplementary Fig. 9.

### Luciferase-based in vitro cytotoxicity assay

For Fig. 4g,h, 2 x 10^4^ Fluc^+^ FUS-CRISPR-engineered Nalm6 cells with 10 min HS (HS) or without (CT) were cultured alone (w/o T), or mixed with αCD19CAR-T cells at an E:T ratio of 1:20 and co-cultured (w/ T) in 96-well plates. Culture media were renewed at 48 h by replacing one-third volume of the supernatant with fresh media. Fluc luminescence was measured 72 h after co-culture using the Bright-Glo Luciferase Assay System (Promega, E2610) and a Tecan Infinite M200 Pro plate reader. Fluc luminescence represents the amount of surviving Nalm6 tumour cells.

Tumour survival (%) of sample X = (Luminescence of X / mean Luminescence of “CT, w/o T” samples) x 100%

Cytotoxicity (%) of CAR-T cells in sample X = 100% - Tumour survival (%) of X

### Animals

Animal studies were approved in Protocol S15285 by UCSD Institutional Animal Care and Use Committee (IACUC). All researchers complied with animal-use guidelines and ethical regulations during animal studies. Six-to-eight weeks old male NOD.Cg-Prkdcscid Il2rgtm1Wjl/SzJ (NSG) mice were purchased from Jackson Laboratory or UCSD Animal Care Program.

### In vivo bioluminescence imaging

In vivo bioluminescence imaging (BLI) of firefly luciferase signals was performed using Lumina LT Series III (PerkinElmer). Firefly luciferase substrate D-luciferin (GoldBio, LUCK-1G) was administered intraperitoneally, followed by BLI approximately 10 min later until capture of the peak signal. Images were analyzed with Living Image software (PerkinElmer). The integrated luminescence reading within a fixed region of interest (ROI) over the tumour was used to represent the tumour size.

### In-house built FUS system

We developed a FUS system with real-time temperature control feedback loop for hyperthermia experiments (Supplementary Fig. 2a-e). A focused 1.1-MHz single element transducer was fabricated in-house using a pre-focused modified PZT (diameter: 70mm, radius of curvature: 65mm, DL-47, Del Piezo Specialties) with a 20 mm hole in the center. A coupling cone (length: 65mm) with an opening (diameter: 4mm) at the tip was 3D-printed and glued to the transducer to hold degassed water through the acoustic path and to guide the ultrasound focus. The opening at the tip of the cone was sealed with acoustically transparent thin-film (Chemplex, 100). Deionized water was degassed with a vacuum pump (Vevor). A function generator (Sanford Research System, SG386) and a 50dB power amplifier (E&I, 325LA) were used to feed pulsed sine waves to the transducer.

For FUS stimulation on cells in vitro (Supplementary Fig. 2b,c), cells were resuspended in 50μl medium in a PCR tube. The cell-containing PCR tube was fixed on the acoustic absorber (Precision Acoustics, F28-SMALL) below the transducer. A needle-type thermocouple (Physitemp Instruments, MT-29/2HT) was inserted into the tube to measure the temperature of the cell medium with a thermometer (Omega, HH806AU). Acoustic gel (Aquasonic, 26354) was applied between the transducer and the tube.

For in vivo FUS stimulation (Supplementary Fig. 2d,e), the anesthetized mouse was placed on its side on the animal bed with an embedded acoustic absorber. The animal bed is placed on a heating plate (Auber Instruments, WSD-30B) set to 37°C to maintain the body temperature of the anesthetized mouse. The needle-type thermocouple was inserted into the tumour region subcutaneously to measure the temperature. Acoustic gel was generously applied. The FUS transducer was placed above the mouse to focus on the tumour.

The temperature readings were fed to a PID controller in real-time to adjust the output power of the function generator to maintain the focal temperature at the target value. All in vivo FUS stimulation was targeted at 43 °C for 10 min or less. The code repository for the PID controller and the device interfaces can be found at https://github.com/phuongho43/ultrasound_pid.

### In vivo tumour model

2 x 10^5^ Nalm6 cells were injected subcutaneously into NSG mice on Day 0. FUS stimulation (43 °C, 10 min) targeted at the tumour region was performed on Day 9 and Day 12 in the FUS+ groups. 2 x 10^6^ CD19CAR-T cells were administered intravenously on Day 10 in the indicated groups. Tumour aggressiveness was monitored by BLI and caliper measurement (volume□=□length□×□width^2^/2).

### Software and statistical analysis

Data were graphed and the corresponding statistical analysis was performed in GraphPad Prism 9.0.0. The detailed statistical test methods were indicated in the corresponding figure legends. Microscopy images were analyzed in Fiji ImageJ2 2.3.0. Schematic figures were created with BioRender.com.

### Reporting summary

Further information on research design is available in the Nature Research Reporting Summary linked to this article.

## Data availability

The main data supporting the results of this study are available within the paper and its Supplementary Information. Other raw data generated during this study are available from the corresponding authors on reasonable request.

## Code availability

The code repository for the PID controller and the device interfaces for the in-house built FUS system can be found at https://github.com/phuongho43/ultrasound_pid.

## Acknowledgements

Y.Wang discloses support for the research described in this study from NIH EB029122, GM140929, HL121365, HD107206, and CA262815. G.K. discloses support for the research described in this study from NIH EB032822.

## Author contributions

Y. Wu, Z.H., Y.L., and Y. Wang conceived and designed the experiments; Y. Wu, Z.H., Y.L., C.Y., Y.S., and Z.Y. performed the experiments; Y. Wu, Y.L., Z.H., and K.S. analyzed the data; C.Y., P.H., L.Z., J.E., Y.Z., and G.K. contributed materials; Y. Wu, Z.H., Y.L., T.L., G.K., S.C., and Y. Wang wrote the paper. All authors reviewed the manuscript and approved the final version.

## Inclusion & ethics statement

All researchers that fulfill authorship criteria have been included in the author list.

## Competing interests

Y. Wang is scientific co-founder and consultant of Cell E&G Inc. and Acoustic Cell Therapy Inc. These financial interests do not affect the design, conduct or reporting of this research. G. Kwong is co-founder of Glympse Bio and Port Therapeutics. This study could affect his personal financial status. The terms of this arrangement have been reviewed and approved by Georgia Tech in accordance with its conflict-of-interest policies. The other authors declare no competing interests.

## Supplementary Information

The supplementary information contains 11 supplementary figures and figure captions, and 5 supplementary tables and table captions.

---

**Supplementary Figure 1.**
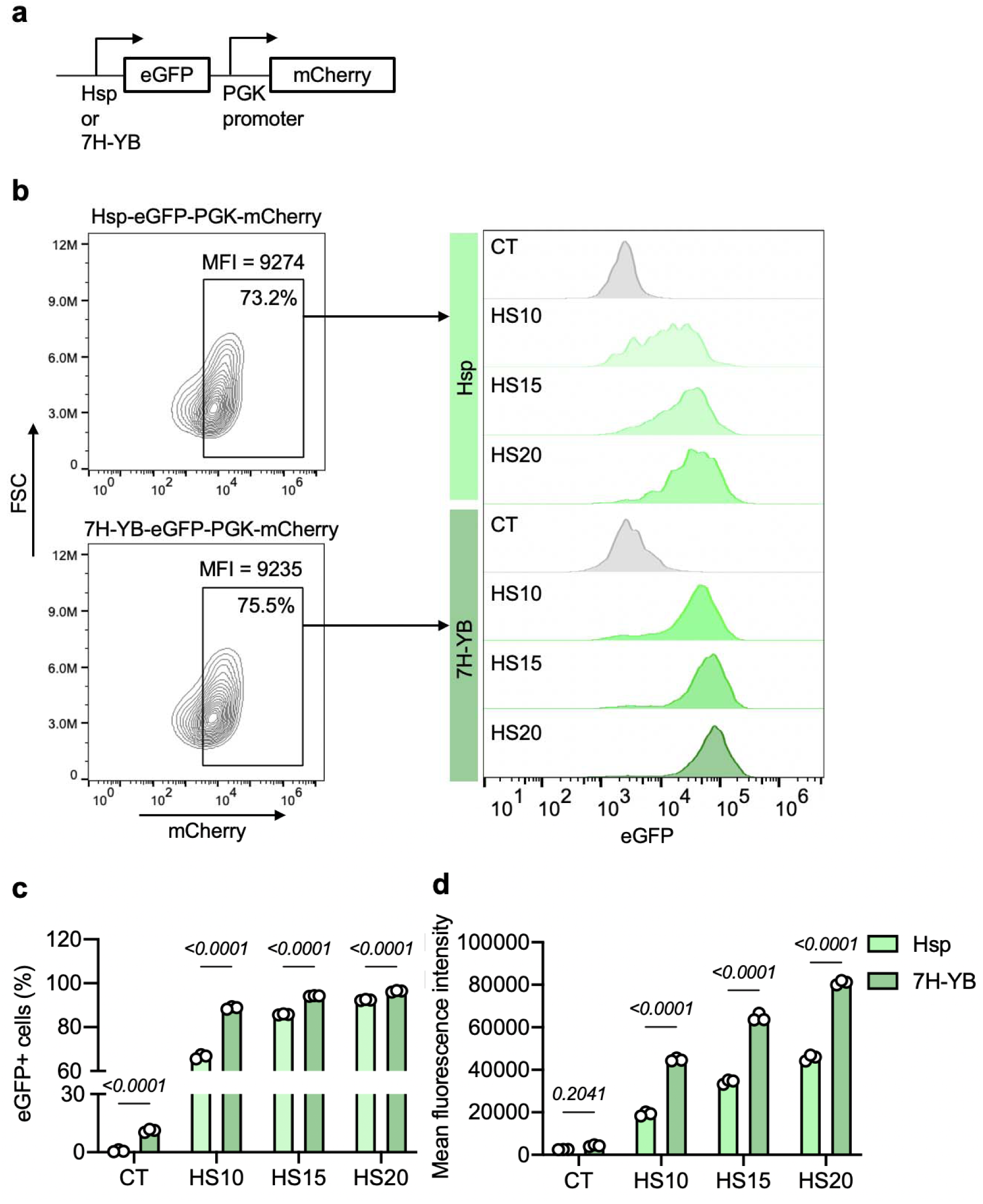
Inducible gene expression controlled by heat-sensitive promoters Hsp and 7H-YB. **a**, Schematics of the Hsp- or 7H-YB-driven eGFP with constitutive mCherry constructs used in this figure. **b**, Representative flow cytometry data of heat-inducible eGFP expression profile in Jurkat cells engineered with Hsp- or 7H-YB-driven constructs in **a**. The same mCherry^+^ cell gate was used in both groups for eGFP expression analysis. **c**,**d**, The percentage of eGFP^+^ cells (**c**) and the mean eGFP fluorescence intensity (**d**) of the above-described engineered Jurkat cells. In **b**-**d**, Cells were treated with no HS (CT), or HS of 10 min (HS10), 15 min (HS15), and 20 min (HS20) and analyzed by flow cytometry 24 h after HS. In **c**,**d**, bar heights represent means; error bars represent s.e.m.; n = 3 technical replicates representative of two independent experiments. Two-way ANOVA followed by Sidak’s multiple comparisons test was used for statistical analysis.

**Supplementary Figure 2.**
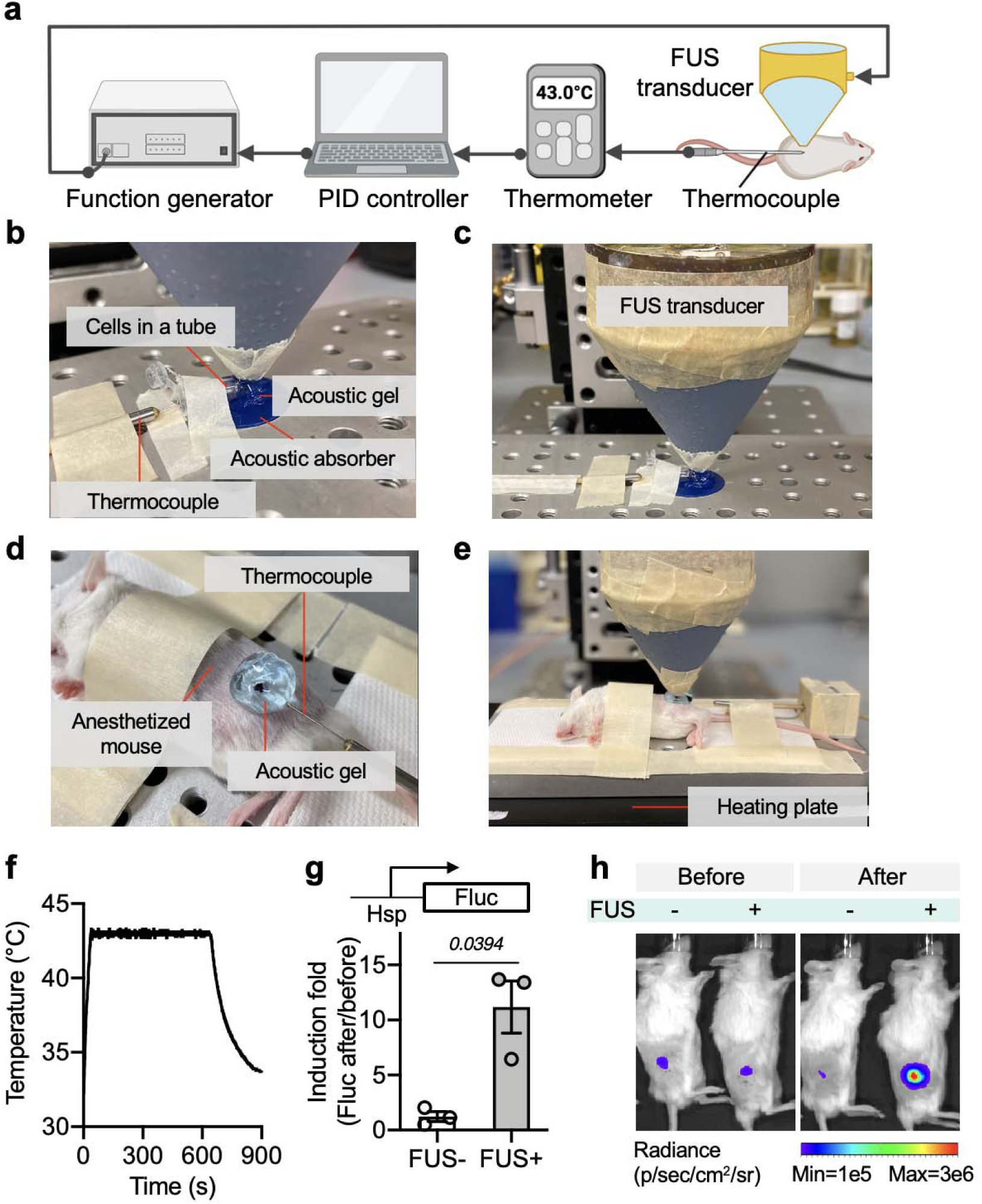
In-house built FUS system. **a**, Schematics of the in-house built FUS system with closed-loop feedback for generation of localized hyperthermia at the target temperature. **b**,**c**, Close-up (**b**) and full shot (**c**) of the experimental setup for FUS stimulation in vitro on cells. **d**,**e**, Close-up (**d**) and full shot (**e**) of the experimental setup for FUS stimulation in vivo. **f**, FUS-induced hyperthermia at 43 °C for 10 min in vivo. **g**,**h**, Quantified induction fold (**g**) and representative images (**h**) of FUS-induced Fluc expression in mice bearing tumors engineered with Hsp-Fluc 6 h after 10 min FUS stimulation at 43 °C. Bar heights represent means; error bars represent s.e.m.; n = 3 mice; paired t test.

**Supplementary Figure 3.**
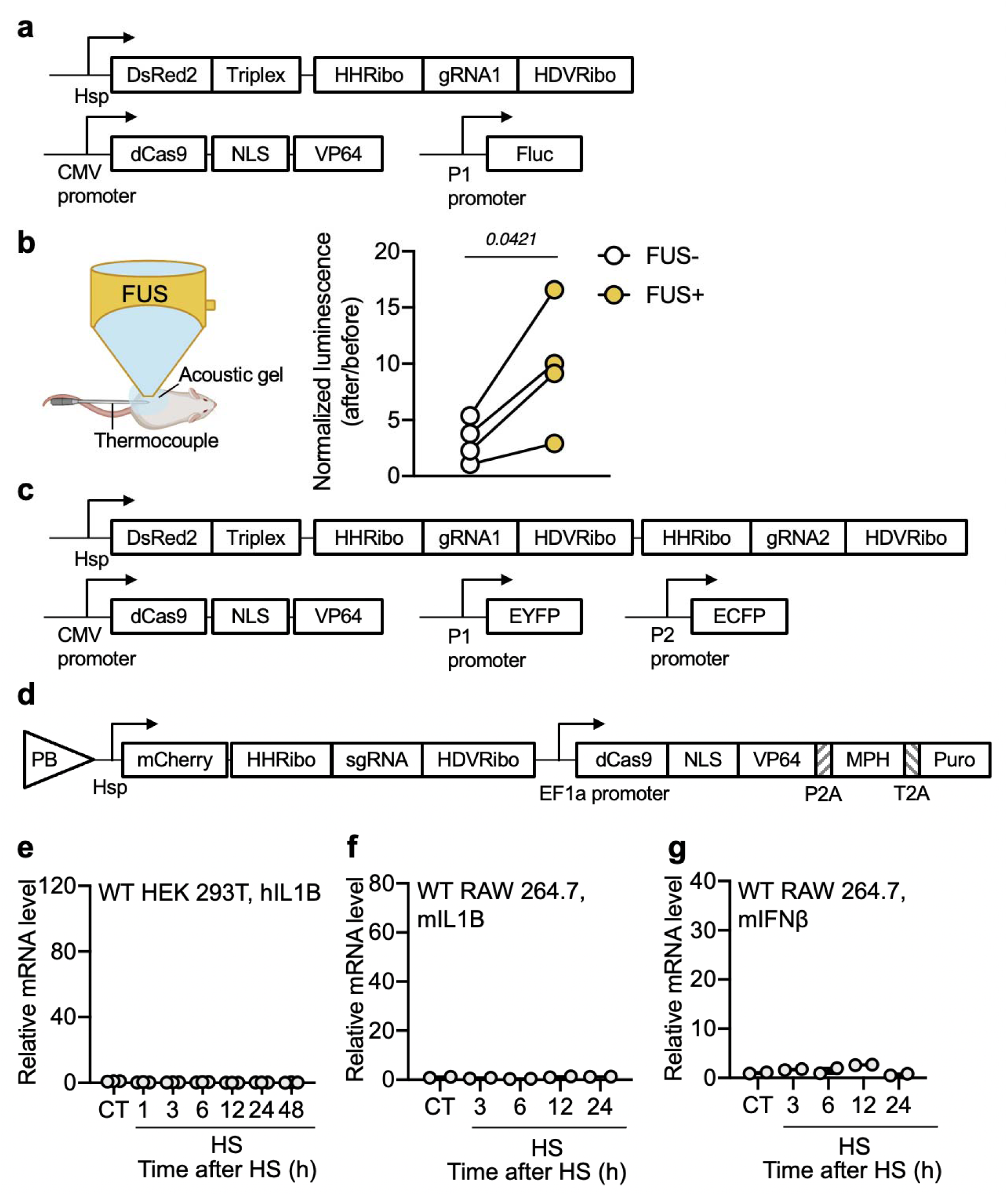
Supplementary figure associated with the FUS-CRISPRa system with the inducible gRNAs. **a**, DNA constructs used in Fig. 1b,c. **b**, In vivo activation of CRISPRa by FUS. Left, schematic illustration of FUS stimulation in vivo. Right, HEK 293T cells transfected with the plasmids in **a** were subcutaneously injected into both sides of NSG mice, followed by FUS stimulation (43 °C, 15 min) 6 h after at one side (FUS+). The other side received no FUS (FUS-). Fluc luminescence of both sides was quantified immediately before and 24 h after FUS stimulation and normalized to the readings before FUS. N = 4 mice. Paired t test. **c**, DNA constructs used in Fig. 1d. **d**, The piggyBac (PB) transposon plasmid used in Fig. 1e-h. Each target gene used a different sgRNA. **e**, Relative hIL1B mRNA levels in non-engineered (wild type, WT) HEK 293T cells without HS (CT), or at different time points after 30 min HS. N = 3 technical repeats. **f**,**g**, Relative mIL1B (**f**) and mIFNβ (**g**) mRNA levels in WT RAW 264.7 cells without HS (CT), or at different time points after 30 min HS. N = 2 technical repeats. Data are representative of two independent experiments.

**Supplementary Figure 4.**
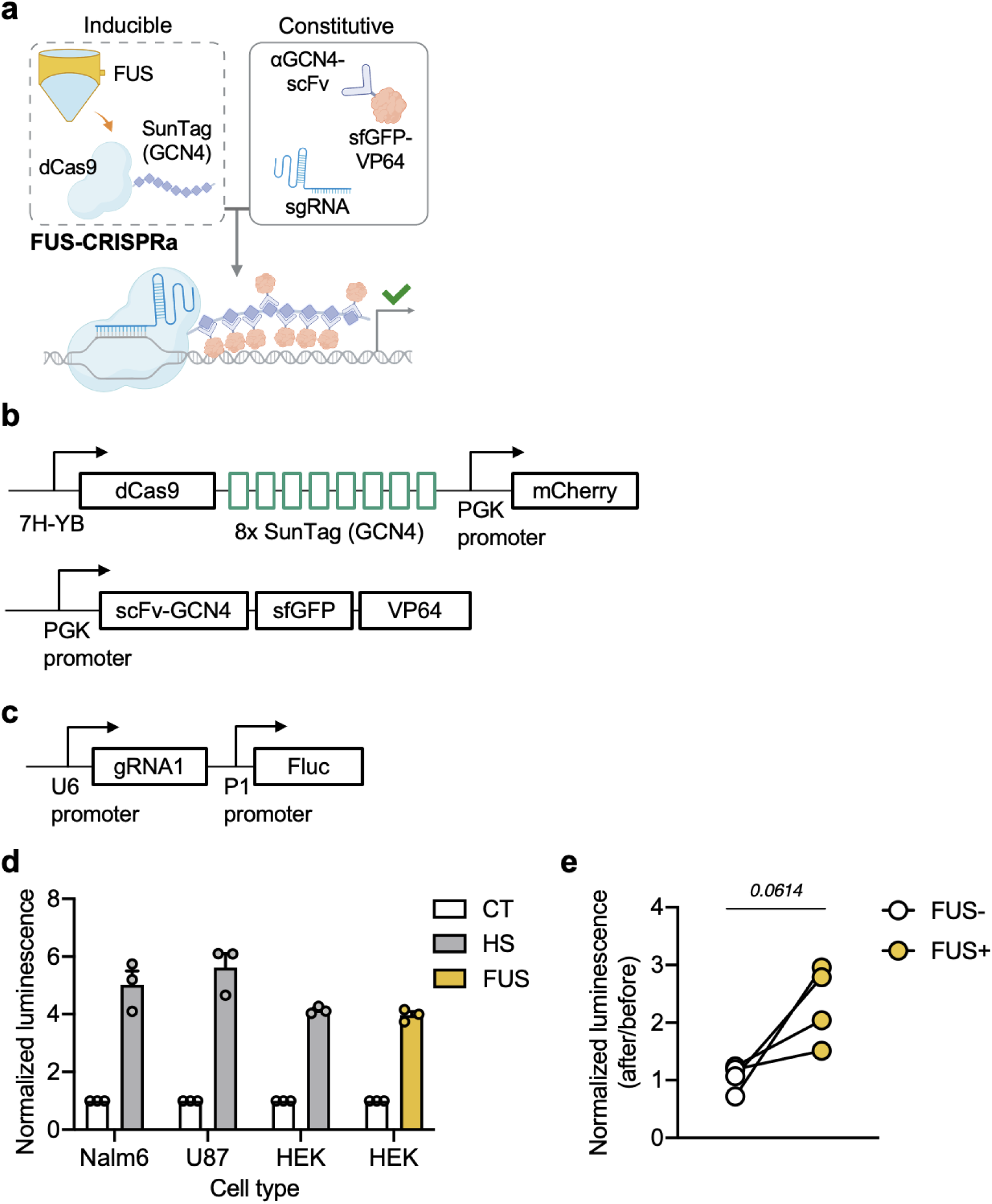
The FUS-CRISPRa system with the inducible dCas9. **a,b,** Schematic illustration (**a**) and DNA constructs (**b**) of the FUS-CRISPRa system with the inducible dCas9 incorporating the SunTag system. **c**, DNA construct containing a constitutive P1-targeting gRNA1 and the P1-driven Fluc. **d**, Normalized Fluc luminescence in multiple cells lines engineered with the lentiviruses encoding the plasmids in **b** and **c**. Readings were quantified 48 h after HS or FUS stimulation and normalized to the corresponding engineered cell lines without HS (CT). HS, with 20 min HS; FUS, 20 min FUS stimulation in vitro on cells. N = 3 biological repeats. **e**, U-87 MG cell line engineered with the P1-targeting FUS-CRISPRa system in **b** and **c** were subcutaneously injected into both sides of NSG mice, followed by FUS stimulation (43 °C, 20 min) 5 days later at one side (FUS+). The other side received no FUS (FUS-). Fluc luminescence of both sides was quantified immediately before and 48 h after FUS stimulation and normalized to the readings before FUS. N = 4 mice. Unpaired t test was used in **d**, paired t test was used in **e**.

**Supplementary Figure 5.**
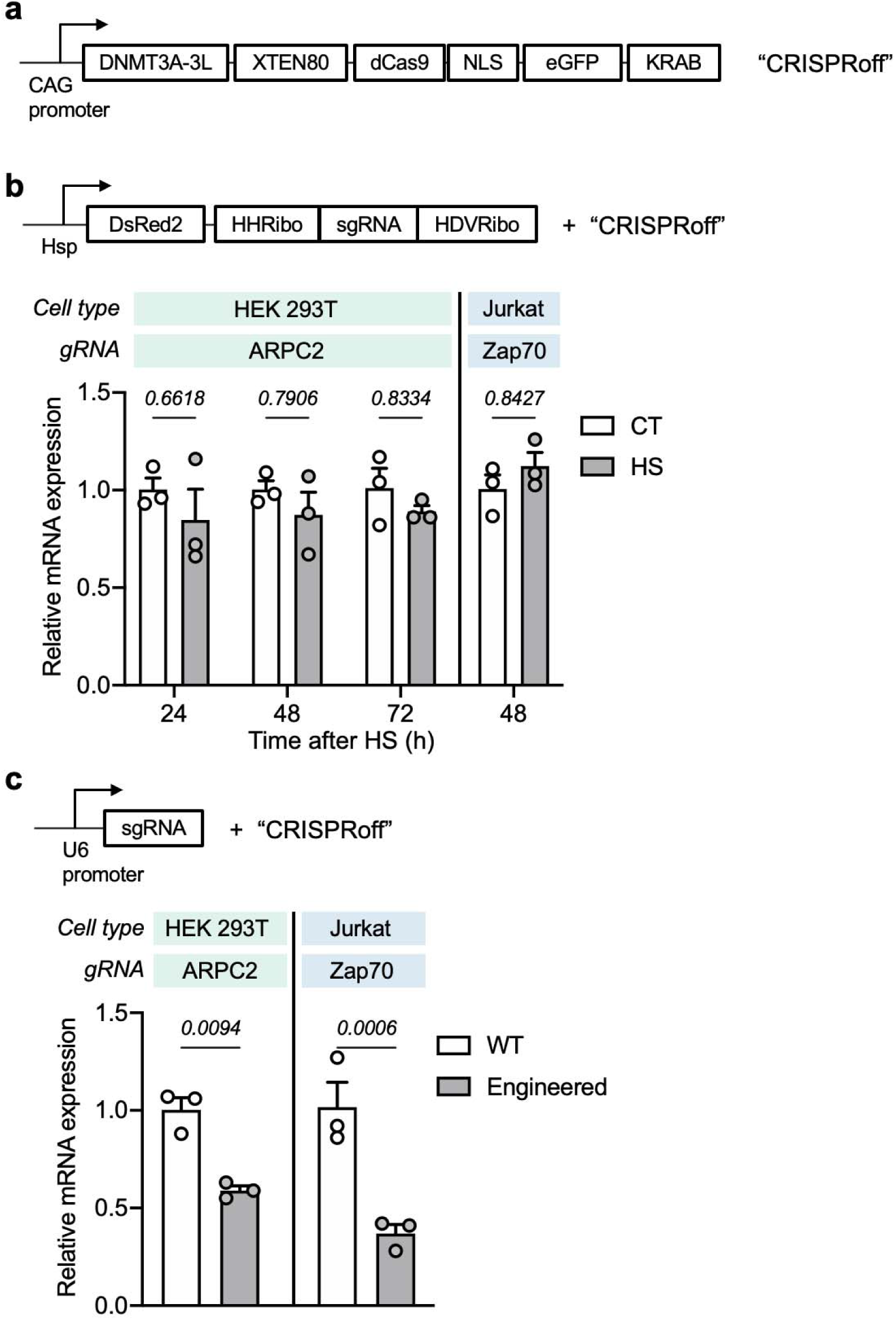
Gene repression with CRISPRoff. **a**, The “CRISPRoff” plasmid used in this figure constructed based on the original CRISPRoff-v2.1 (Addgene plasmid #167981)^46^. **b**, Relative mRNA expression of target genes in different cell types engineered with Hsp-RGR and CRISPRoff. **c**, Relative mRNA expression of target genes in different cell types engineered with constitutive gRNA and CRISPRoff three days after transfection. In **b**,**c**, bar heights represent means; error bars represent s.e.m.; n = 3 technical repeats. Data are representative of two independent experiments. Two-way ANOVA followed by Sidak’s multiple comparisons test was used for statistical analysis.

**Supplementary Figure 6.**
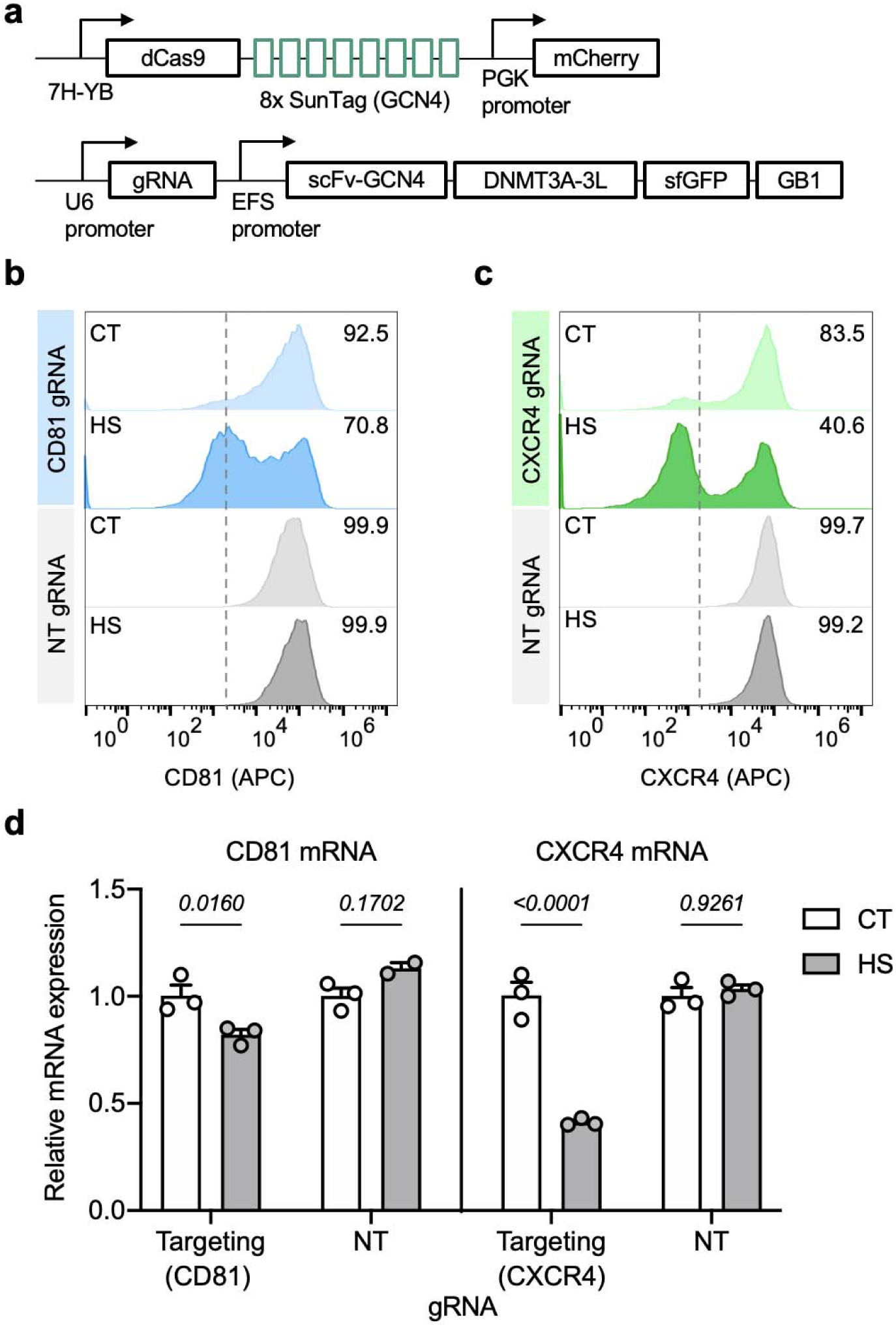
Gene repression in Nalm6 cells engineered with FUS-CRISPRi targeting CD81 or CXCR4. **a**, The FUS-CRISPRi constructs. **b**,**c**, Representative staining results of CD81 (**b**) or CXCR4 (**c**) in the engineered Nalm6 cells four days after HS. **d**, Relative CD81 and CXCR4 mRNA expression in cells in **b** and **c** quantified three days after HS. HS, with 15 min of HS; CT, without HS. In **d**, bar heights represent means; error bars represent s.e.m.; n = 3 technical replicates. Data are representative of two individual experiments. Two-way ANOVA followed by Sidak’s multiple comparisons test was used for statistical analysis.

**Supplementary Figure 7.**
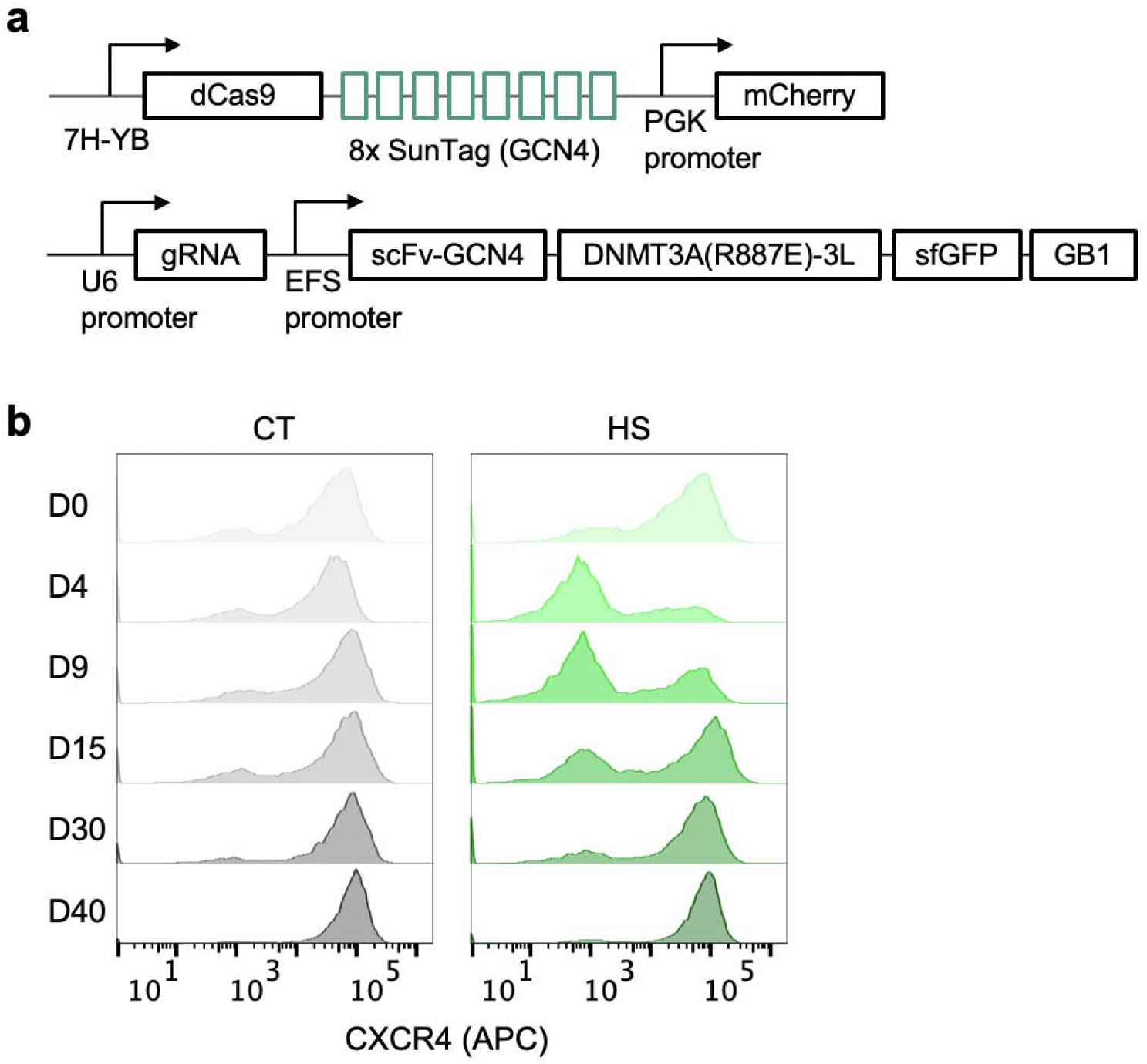
Reversible gene repression via FUS-CRISPRi. **a**, Schematics of FUS-CRISPRi using the R887E mutant DNMT. **b**, Flow cytometry profile of CXCR4 staining in cells engineered with FUS-CRISPRi targeting CXCR4 at different time points after HS. HS, with 20 min HS; CT, without HS.

**Supplementary Figure 8.**
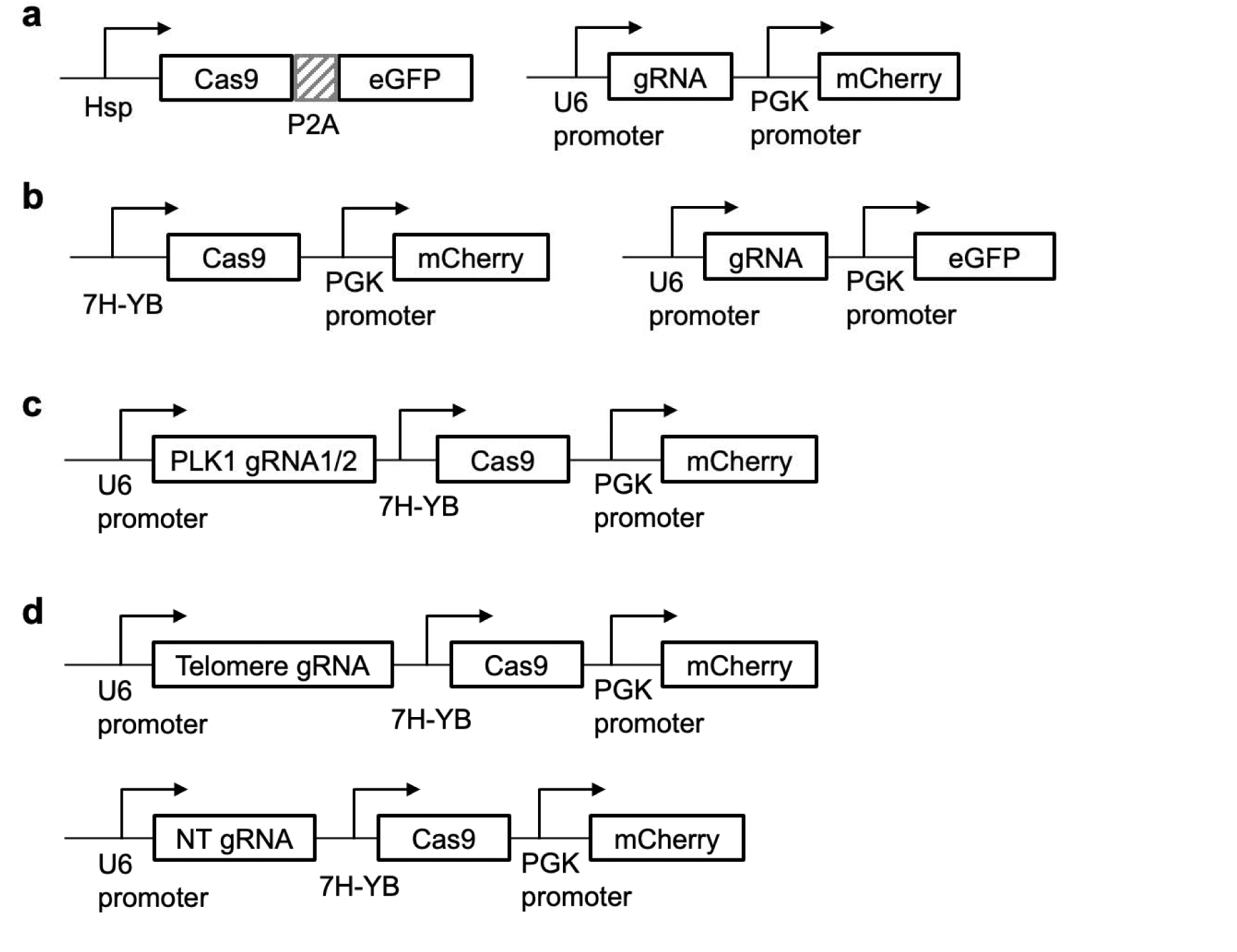
Constructs used in the FUS-CRISPR system. **a**,**b**, “Two-plasmid” design of the FUS-CRISPR system with Hsp (**a**) or 7H-YB (**b**) and different arrangement of marker fluorescent proteins. **c**, The “all-in-one” construct used for FUS-CRISPR targeting PLK1 gene. **d**, DNA constructs for FUS-CRISPR with telomere-targeting gRNA or NT gRNA.

**Supplementary Figure 9.**
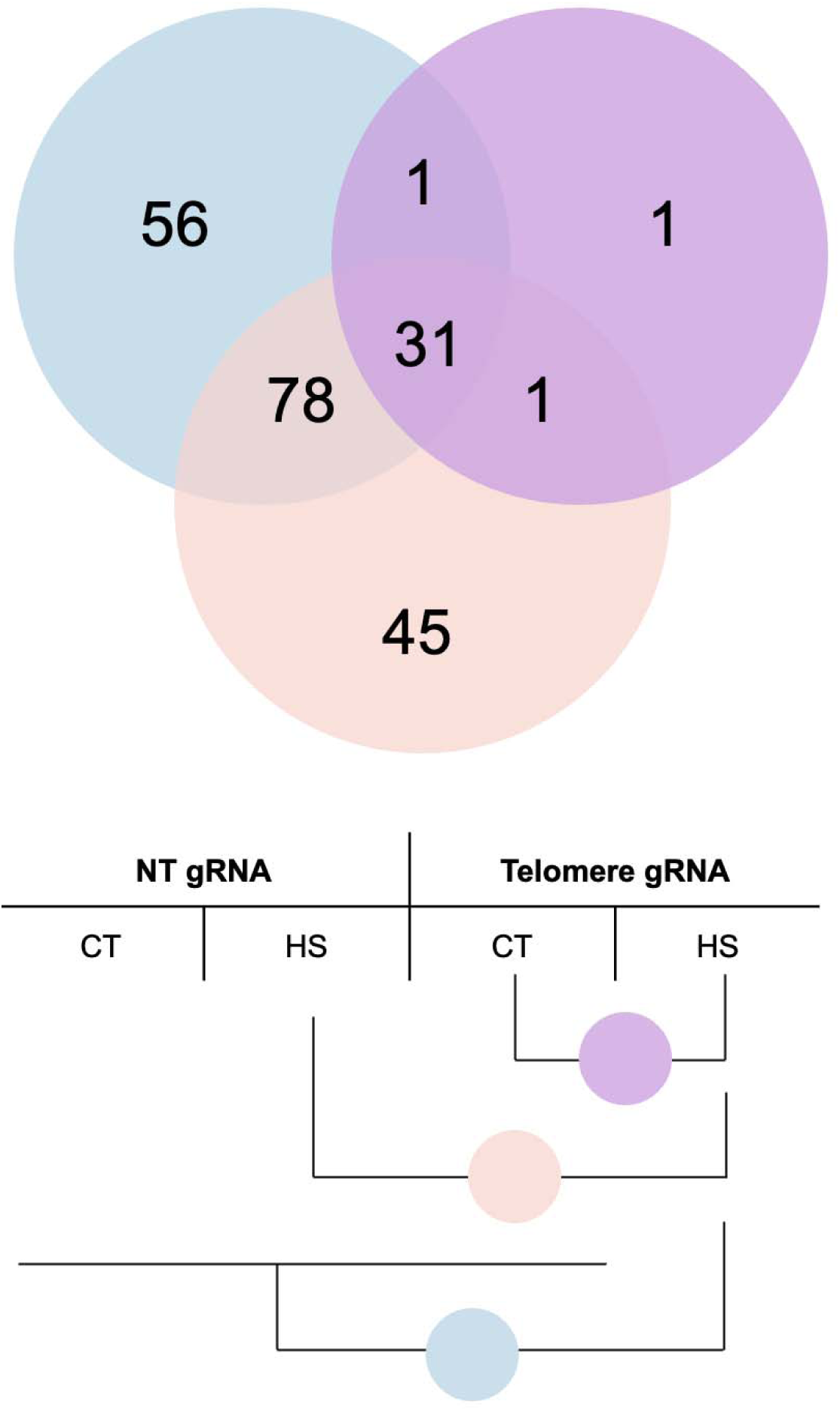
Venn diagram summarizing the differentially expressed genes in the illustrated three groups of comparisons from the RNA-seq data.

**Supplementary Figure 10.**
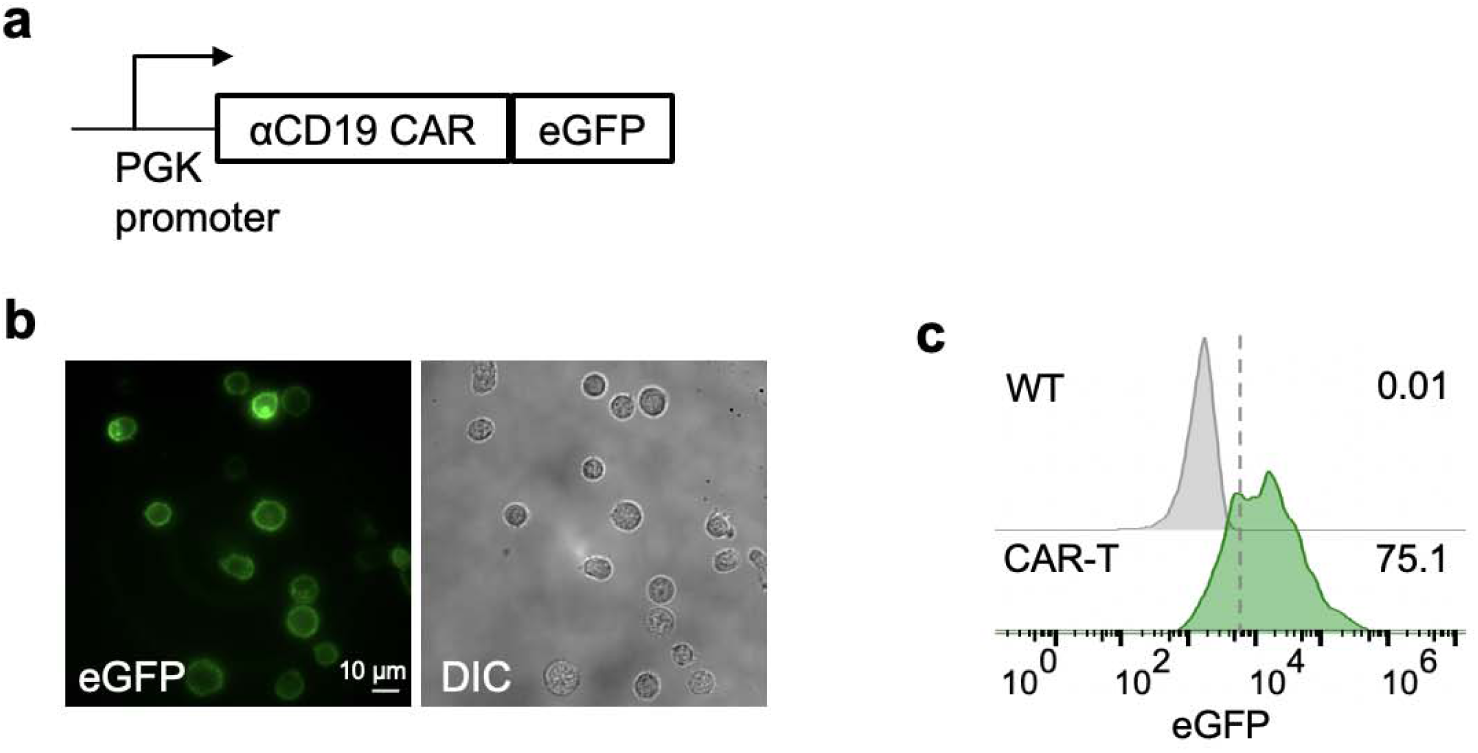
Anti-CD19 CAR-T cells. **a**, The anti-CD19 (αCD19) CAR plasmid. **b**, Primary human T cells expressing the construct in **a** with membrane localization of eGFP. Scale bar = 10 μm. **c**, Representative eGFP expression profiles of WT primary human T cells and the αCD19CAR-T cells in **b**.

**Supplementary Figure 11.**
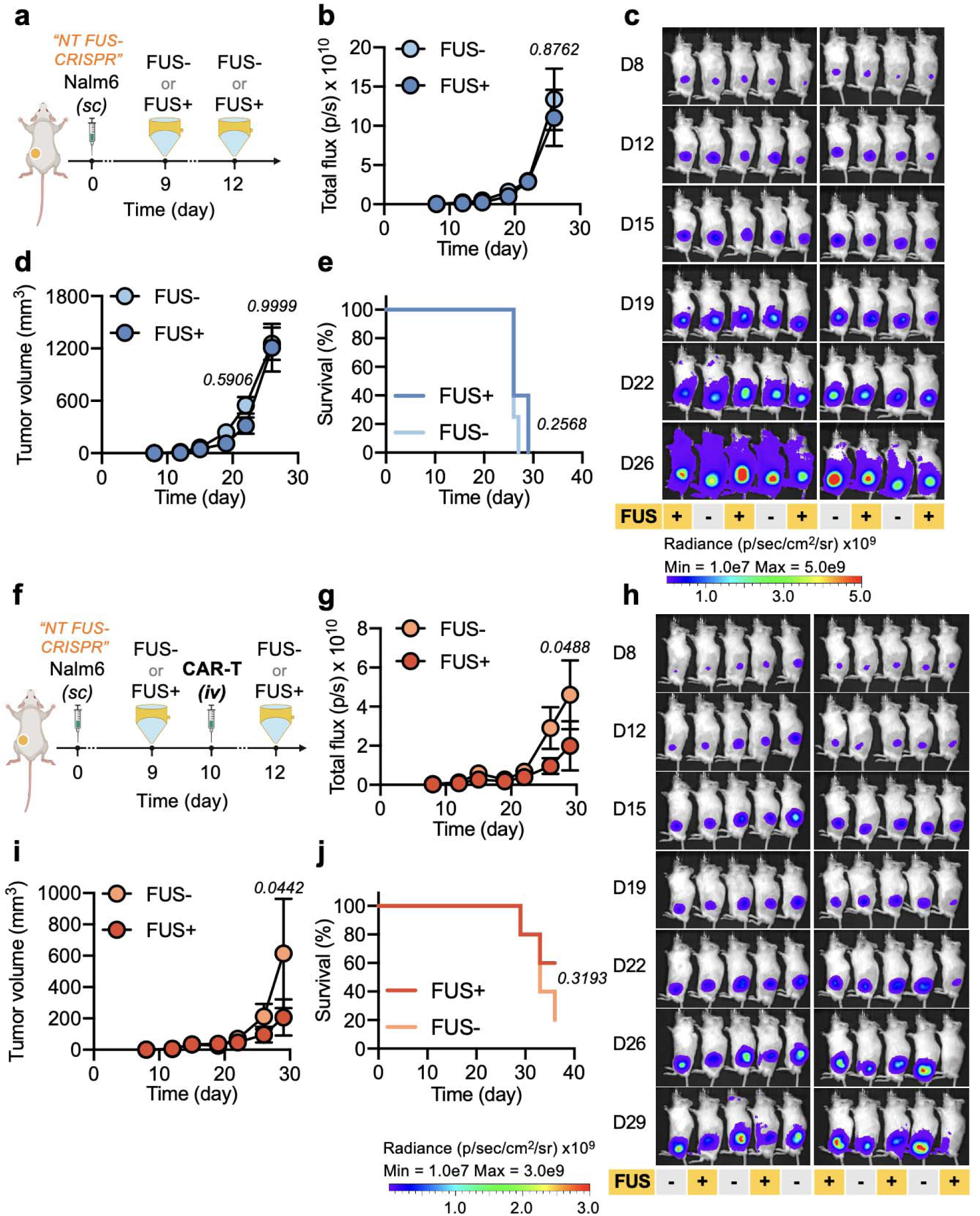
Control experiments related to Figure 5 using FUS-CRISPR with a non-targeting gRNA. **a**, Timeline of experiment in NSG mice. **b-d**, Tumor aggressiveness in the mice in **a** quantified by total flux of the tumor from BLI measurement (**b**), the corresponding BLI images (**c**), and the tumor volume based on caliper measurement (**d**). **e**, Survival curves of the tumor-bearing mice in **a**. **f**, Experimental timeline of FUS-CRISPR with NT gRNA combined with CAR-T therapy in NSG mice. **g-i**, Tumor aggressiveness in the mice in **f** quantified by total flux of the tumor (**g**), the corresponding BLI images (**h**), and the caliper-measured tumor volume (**i**). **j**, Survival curves of the tumor-bearing mice in **f**. Data points represent means; error bands represent s.e.m.; n = 5 mice per group. Two-way ANOVA followed by Sidak’s multiple comparisons test was used in **b**, **d**, **g**, and **i**. Log-rank (Mantel-Cox) test was used in **e** and **j**.

**Supplementary Table 1.**
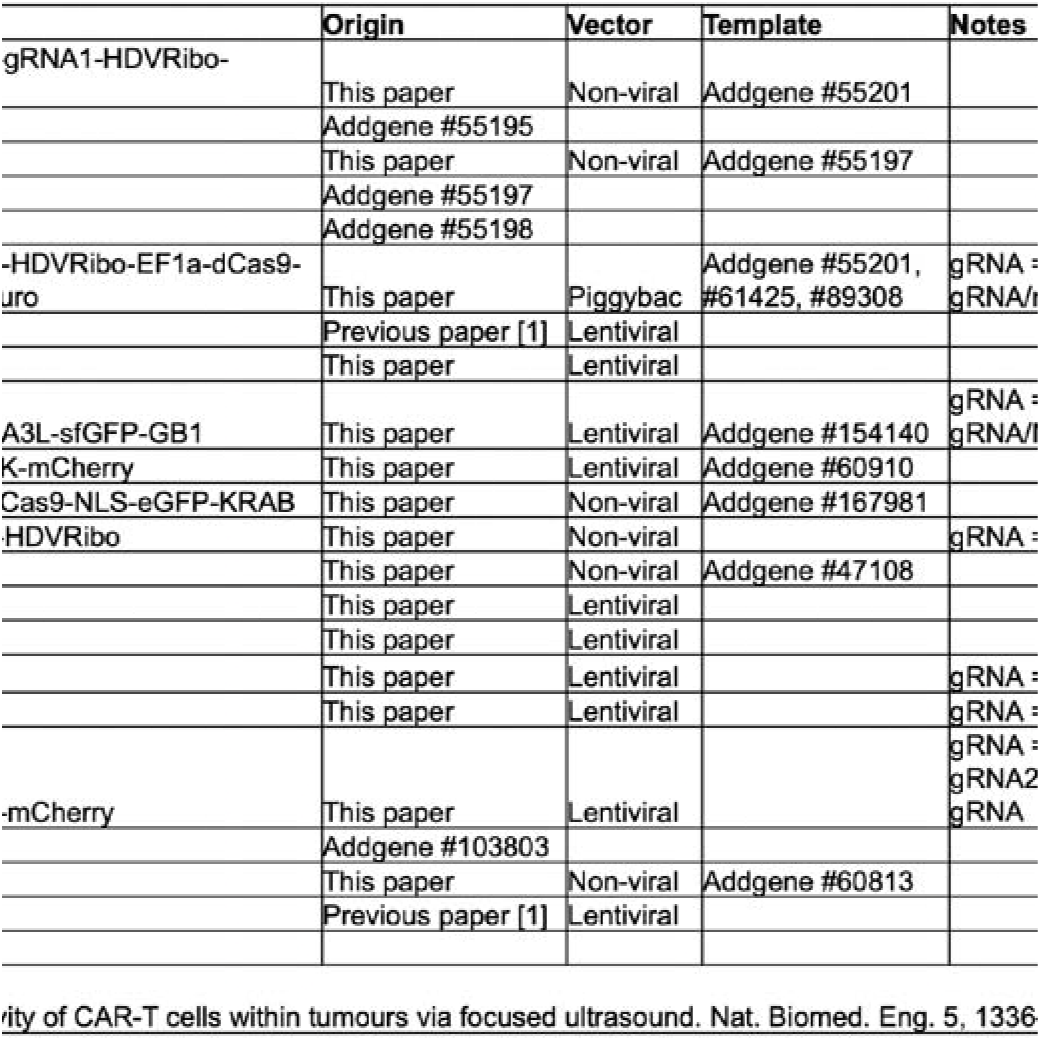
List of plasmids used in this study.

**Supplementary Table 2.**
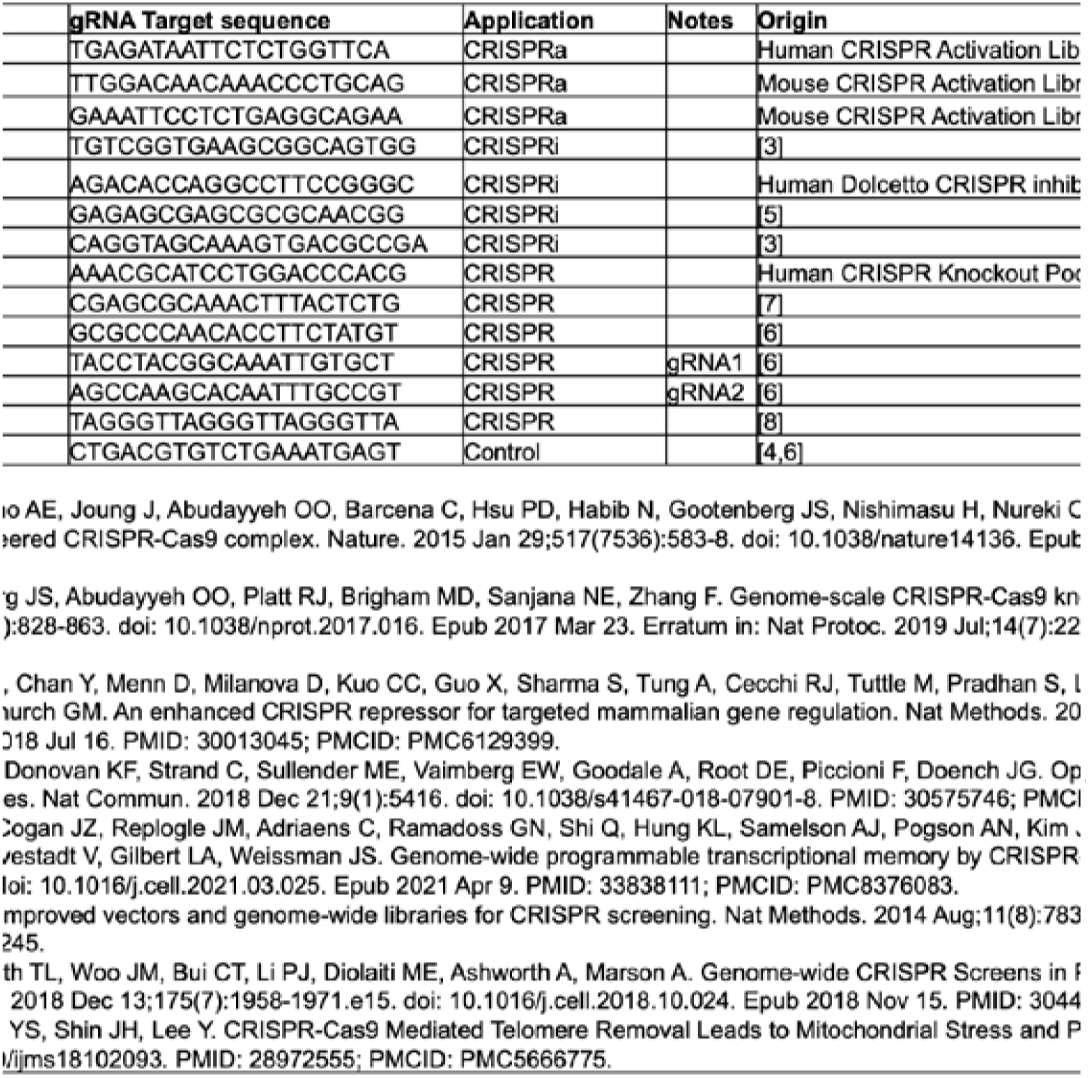
Sequences of gRNAs used in this study.

**Supplementary Table 3.** List of antibodies used in this study.

| ID | Antibody | Company | Catalog number | Application |
| --- | --- | --- | --- | --- |
| 1 | Anti-IL-1 beta antibody | Abcam | ab2105 | Western blot |
| 2 | β-actin | Santa Cruz | sc-69879 | Western blot |
| 3 | APC anti-human CD81 (TAPA-1) | Biolegend | 349509 | Flow cytometry |
| 4 | APC anti-human CD184 (CXCR4) | Biolegend | 306509 | Flow cytometry |
| 5 | APC anti-human CD69 | Biolegend | 310910 | Flow cytometry |
| 6 | Anti-T-Cell Receptor Antibody, clone C305 | Sigma-Aldrich | 05-919 | TCR activation |
| 7 | Janelia Fluor® HaloTag® Ligands | Promega | GA1120 | Fluorescence imaging |

**Supplementary Table 4.**
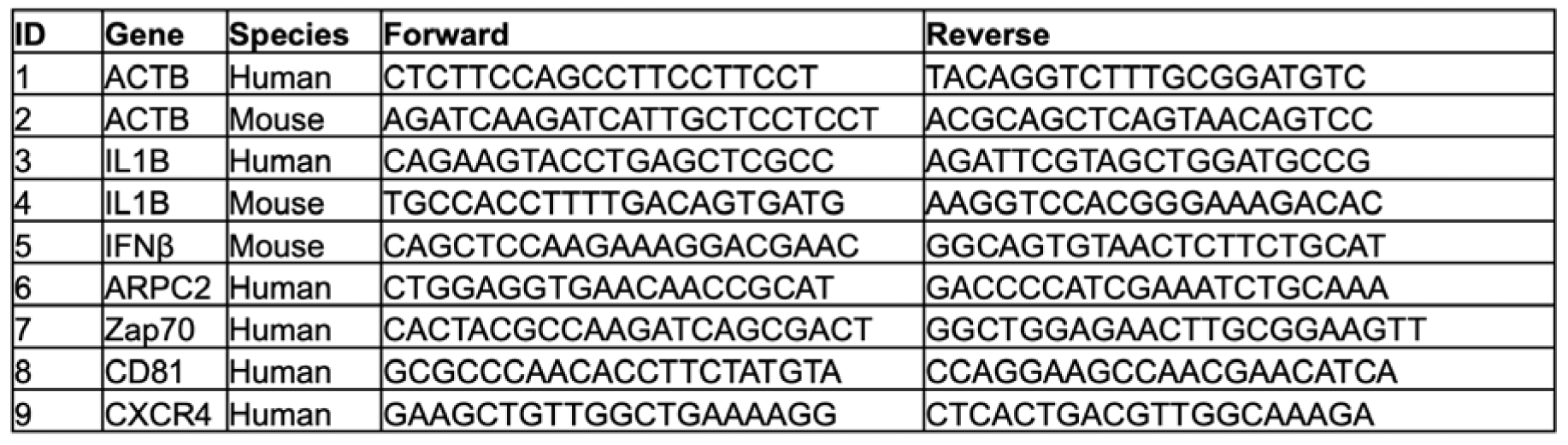
Sequences of qPCR primers used in this study.

**Supplementary Table 5.**
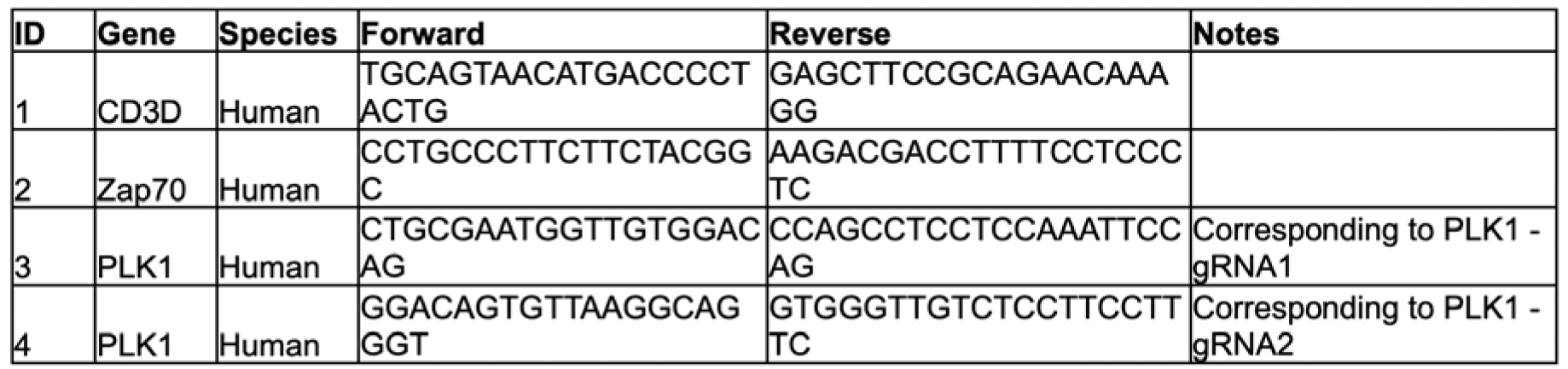
Sequences of genotyping PCR primers used in this study.

